# Repeated morphine reorganizes sleep-wake states, cortical and central medial thalamic oscillations, and network synchronization in mice

**DOI:** 10.64898/2026.09.20.752991

**Authors:** Vasilije Tadic, Mackenzie Walz, Slobodan M Todorovic, Tamara Timic Stamenic

## Abstract

**Objectives:** Opioids disrupt sleep, but how repeated exposure reorganizes thalamocortical networks is unknown. We asked whether morphine alters sleep architecture, regional oscillations, and phase synchronization, and how exposure history modifies these effects.

**Methods:** From mice, we analyzed cortical electroencephalogram (EEG), central medial thalamic (CMT) local field potentials (LFP), and electromyography (EMG) for 24h after the first and fourth injections of morphine and in abstinence, and quantified vigilance-state occupancy, spectral power, state-power correlations, and the weighted phase-lag index (wPLI), which minimizes volume conduction.

**Results:** Morphine produced hyperlocomotion, wake promotion persisting with repeated exposure, non-rapid eye movement (NREM) sleep suppression, complete rapid eye movement (REM) sleep loss, and rebound sleep. Cortex and thalamus dissociated: in NREM sleep after the first injection, cortical delta and low-gamma power increased, while CMT delta, theta, alpha, and beta power decreased. NREM occupancy coupled more strongly to CMT delta and theta power after both injections, whereas cortical wake correlations in theta, alpha, and beta fell and reversed during abstinence. Overall synchronization was altered only in theta and alpha: corticocortical alpha synchronization decreased acutely, and repeated exposure reduced thalamocortical theta synchronization in wake and NREM sleep. During abstinence, sleep architecture and synchronization largely normalized while spectral and state-power abnormalities persisted alongside mechanical hypersensitivity.

**Conclusions:** We found that morphine does not merely reduce sleep; it reorganizes it: cortex and CMT move in opposite directions, thalamocortical phase synchronization is redistributed, and the resulting spectral and state-power abnormalities outlast the recovery of sleep architecture itself rather than producing tolerance.

**Support:** NIH grants: R35GM141802, K01DA055258

## INTRODUCTION

Opioid use is strongly associated with disrupted sleep, and sleep abnormalities are increasingly recognized as an important component of opioid use disorder and opioid withdrawal^1^. Opioids alter sleep architecture by increasing wakefulness, reducing non-rapid eye movement (NREM) sleep, and suppressing rapid eye movement (REM) sleep; these disturbances extend beyond the period of acute drug exposure^1,2^. Rodent models of opioid exposure reproduce many of these disruptions and allow the underlying neural mechanisms to be examined directly. In various experimental models, morphine produces pronounced wake promotion and suppression of NREM and REM sleep, indicating that opioid-induced sleep disruption is a direct neurobiological consequence of opioid exposure rather than solely a consequence of disease-related or environmental factors^3,4^. One established mechanism is engagement of μ-opioid receptors (MORs) in the ventrolateral preoptic area (VLPO), a critical sleep-promoting region: in rats, an acute dose of morphine promotes wakefulness by activating VLPO MORs, thereby inhibiting VLPO neurons and suppressing sleep-promoting mechanisms^5^.

It is well known that the thalamus is a key node in arousal and sleep regulation, in particular, the midline nuclei such as the paraventricular nucleus (PVT) and the intralaminar nuclei such as the central medial nucleus (CMT)^6–9^. These regions interact with sleep-wake circuits and reward pathways and are affected by opioid exposure in ways that can substantially alter sleep^10–12^. The midline and intralaminar thalamic complex are a well-documented component of the brainstem-diencephalic-thalamocortical circuitry that regulates affective, cognitive, and executive function^13^, and the dense innervation of the rostral intralaminar complex by the reticular formation places it within the ascending reticular activating system (reviewed in^14^).

Within this system, the PVT is the best-characterized opioid-sensitive component. PVT neurons are wake-active and causally promote transitions from sleep to wakefulness^9^, and the nucleus is anatomically positioned to integrate circadian, hypothalamic, and brainstem arousal signals^12,15^. MOR-expressing PVT neurons have been directly implicated in morphine-induced wakefulness, establishing the PVT as an opioid-sensitive substrate for sleep-wake disruption^10^. The CMT, by contrast, has been strongly implicated in the regulation of NREM sleep rather than in arousal alone^6,7^. Both tonic and burst firing of CMT neurons modulate cortical activity during sleep, providing dual control over sleep-wake states^7^, and the CMT appears to function as a critical hub through which both general anesthesia and natural sleep are initiated^6^. It has been reported that morphine engages this nucleus: acute morphine administration (10 mg/kg) induces c-Fos expression in the midline PVT, and the intralaminar CMT^16^, and morphine-induced conditioned place preference training elevates c-Fos expression specifically in the CMT^17^. Despite this evidence and the CMT’s established roles in sleep-wake transitions, cortical slow-wave synchronization, and the homeostatic response to prolonged wakefulness, considerably less is known about how morphine affects the CMT than about its effects on the PVT. This gap matters because the CMT not only influences arousal but also organizes NREM slow waves, both under baseline conditions and during sleep recovery after extended wakefulness.

To our knowledge, the effects of repeated morphine exposure on CMT oscillatory activity and on corticocortical and thalamocortical phase synchronization across vigilance states have not been characterized previously. We therefore examined how repeated morphine exposure and subsequent abstinence affect sleep-wake architecture and oscillatory activity in the CMT and barrel cortex, with the aim of determining how the acute wake-promoting effects of morphine interact with sleep-homeostatic mechanisms across repeated exposure and abstinence. Because CMT activity is functionally linked to cortical state regulation, local CMT oscillations alone may not capture opioid-induced changes in this system. So, we assessed both frequency-specific power within the CMT and functional phase synchronization between the CMT and the barrel cortex, quantified with the weighted phase lag index (wPLI), a measure that weights each phase-difference estimate by its magnitude and is therefore comparatively insensitive to zero-lag coupling arising from volume conduction and shared sources^18^. This approach allowed us to determine whether morphine alters local thalamic oscillatory activity, the coordination of thalamocortical networks across wakefulness and NREM sleep, or both. Finally, because CMT activity has been implicated in the homeostatic response to prolonged wakefulness, we examined whether morphine altered the relationship between the time spent in wakefulness or NREM sleep and cortical and thalamic spectral power.

## MATERIALS AND METHODS

### Animals

Adult male and female C57BL/6J mice (8-16 weeks of age, The Jackson Laboratory, Bar Harbor, ME, USA) were used in this study. Animals were housed in a temperature- and humidity-controlled vivarium under a 14:10h light-dark cycle with ad libitum access to food and water (lights-on at 6 am, Zeitgeber time (ZT0) and lights off at 8 pm (ZT14)). Mice were group-housed whenever possible and monitored daily for health and welfare. All experimental procedures were conducted in accordance with the National Institutes of Health “Guide for the Care and Use of Laboratory Animals” and were approved by the Institutional Animal Care and Use Committee (IACUC) of the University of Colorado Anschutz Medical Campus. Every effort was made to minimize animal suffering and to reduce the number of animals used.

### Morphine administration and sleep experimental design

Morphine sulfate (4 mg/mL; Fresenius Kabi USA) was administered intraperitoneally (i.p.) at 15 mg/kg. Control animals received an equivalent volume of saline (Fresenius Kabi USA) by the same route. The animals were given saline during the first week and, after a recovery period, received morphine (N=7, within-subjects design; Figure 1a). Morphine or saline was administered at ZT9 on four consecutive days (Figure 1a). Continuous EEG/LFP, EMG, and time-locked video recordings were analyzed for 24h after the first injection, for 24h after the fourth injection, and for 24h during abstinence (after the final injection). The 1h-bin analysis is reported together across 24h recording period, with the average value of episode length (mean episode duration), stage length (cumulative time in each state), as well as the number of episodes for all three intervals, as indicated in the individual experiments and figures: the acute effect (the first 5h after injection, which falls entirely within the light phase and ends at lights-off), the dark phase (10h), and the subsequent light phase (9h).

**Figure 1.**
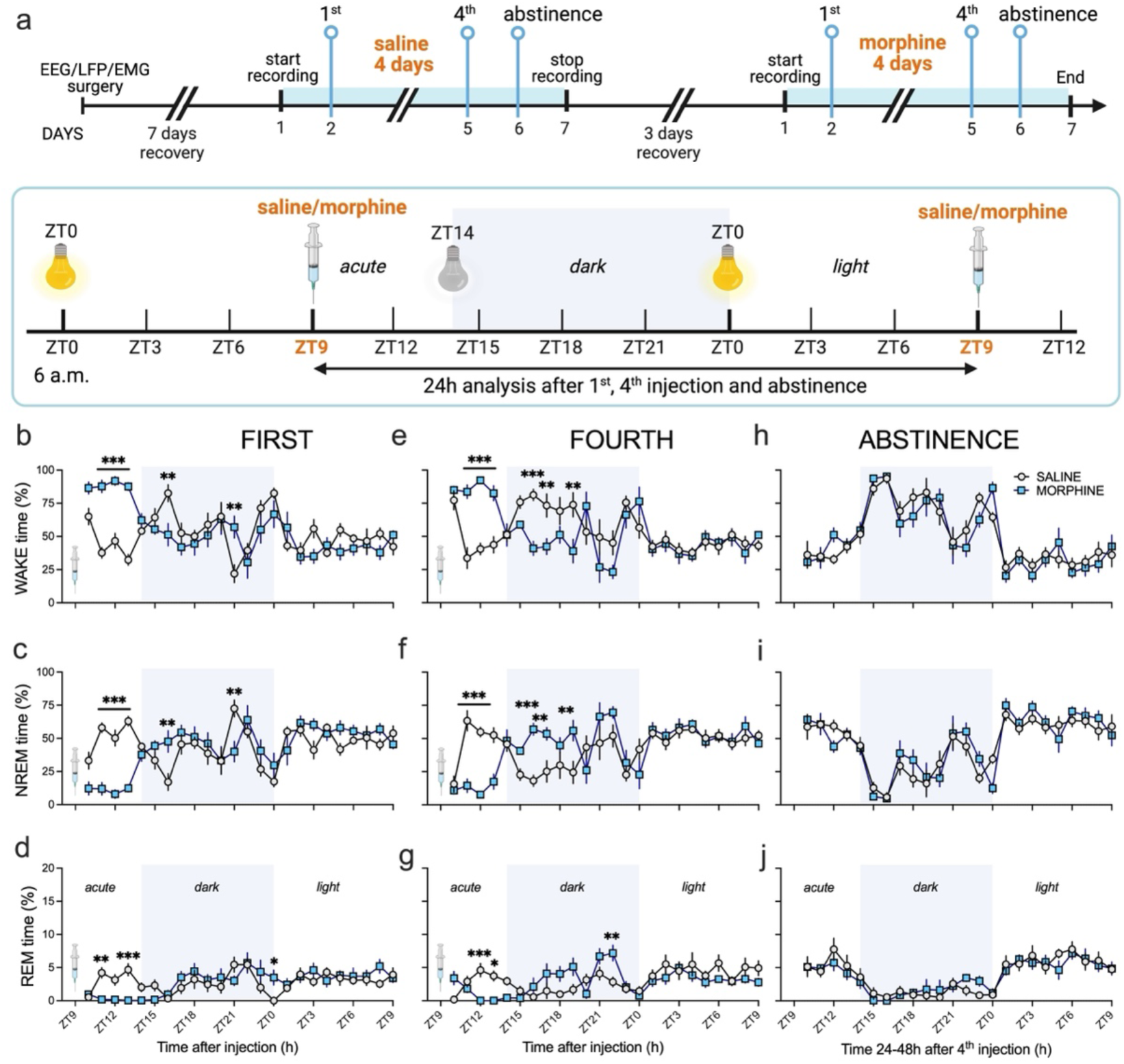
Effects of repeated morphine administration and abstinence on vigilance states across the light–dark cycle. (a) Experimental design and recording schedule. Rats were implanted with EEG/LFP/EMG electrodes and allowed to recover before recordings. Animals received daily injections of saline or morphine at ZT9 for four consecutive days. Continuous recordings were analyzed during the 24h following the first injection, the fourth injection, and after morphine abstinence. The light–dark cycle consisted of 14h light and 10h dark. Shaded regions indicate the dark phase. (b-d) Time course following the <u>first injection</u> of (b) <u>wakefulness</u> (wake time, %): two-way RM ANOVA: Time F_23,138_=5.99, p<0.0001; Morphine F_1,6_=2.92, p=0.138; Interaction F_23,138_=7.29, p<0.0001, Šídák’s post hoc presented in the figure; (c) <u>NREM sleep</u>; two-way RM ANOVA: Time F_23,138_=6.06, p<0.0001; Morphine F_1,6_=2.19, p=0.189; Interaction F_23,138_=7.34, p<0.0001, Šídák’s post hoc presented in the figure; and (d) <u>REM sleep</u>: two-way RM ANOVA: Time F_23,138_=4.05, p<0.0001; Morphine F_1,6_=0.04, p=0.8392; Interaction F_23,138_=3.62, p<0.0001, Šídák’s post hoc presented in the figure. (e–g) Time course following the <u>fourth injection</u> of (e) <u>wakefulness</u>: two-way RM ANOVA: Time F_23,138_=6.69, p<0.0001; Morphine F_1,6_=0.05, p=0.832; Interaction F_23,138_=8.01, p<0.0001, Šídák’s post hoc presented in the figure; (f) <u>NREM sleep</u>; two-way RM ANOVA: Time F_23,138_=7.06, p<0.0001; Morphine F_1,6_=0.20, p=0.670; Interaction F_23,138_=8.21, p<0.0001, Šídák’s post hoc presented in the figure; and (g) <u>REM sleep</u>; two-way RM ANOVA: Time F_23,138_=3.85, p<0.0001; Morphine F_1,6_=4.46, p=0.08; Interaction F_23,138_=4.58, p<0.0001, Šídák’s post hoc presented in the figure. (h–j) Time course following abstinence of wakefulness (h), NREM sleep (i), and REM sleep (j); statistical analysis was not statistically significant. Data are presented as mean ± SEM, N=7 animals. Open circles represent the saline group, and blue squares represent the morphine group. The syringe symbol indicates the injection time (ZT9). Asterisks denote significant differences between saline- and morphine-treated animals at the corresponding time points - Šídák’s. Shaded areas indicate the dark period. ZT, Zeitgeber time.

### Locomotor activity

Locomotor activity was assessed in an open arena (22×22×28 cm). Mice were placed individually in the center of the arena and allowed to explore freely for 1h immediately after receiving morphine (15 mg/kg) or saline i.p. on four consecutive days (N=8 animals per group, between-subject design). Movement was recorded with an overhead video camera and analyzed with ANY-maze software (Stoelting Co., Wood Dale, IL, USA). Distance traveled, mean locomotor speed, and time immobile were quantified across the full 1h test session on each of the four days; immobility was defined as movement below 2 cm/s. The arena was thoroughly cleaned with 70% ethanol between animals to remove olfactory cues, and all testing was performed under consistent lighting and environmental conditions. Behavioral data were collected and analyzed by investigators blinded to group assignment. Locomotor measures were compared between saline- and morphine-treated groups.

### Mechanical Sensitivity Assessment (von Frey Test) during abstinence

Mechanical sensitivity was assessed using an electronic von Frey apparatus (Ugo Basile, Varese, Italy) as previously described^19^. Animals were placed individually in clear plastic enclosures on an elevated wire-mesh platform and allowed to acclimate for 30 min before testing. A single rigid filament tip was then applied to the mid-plantar surface of the hind paw, and the device applied an increasing upward force (0-50 g). Force application ceased on paw withdrawal, and the maximum force eliciting a sharp flinch or lifting response was recorded electronically as the paw withdrawal threshold (PWT) in grams. Baseline measurements were obtained before the first morphine or saline injection and repeated 24h after the last injection, at the beginning of abstinence (N=9 per group, between-subject design). Both hind paws were tested at least 3 times, and the average withdrawal threshold was used in the statistical analysis. Assessments were performed by an experimenter blinded to group assignment.

### EEG/EMG Surgery and Electrophysiological Recordings

Surgical implantation and electrophysiological recordings followed procedures described previously^20–22^. Mice were anesthetized with isoflurane (5% for induction and 1.5-3% for maintenance) and placed in a stereotaxic frame for chronic implantation of cortical, thalamic, and electromyographic electrodes. A coated tungsten electrode was implanted into the CMT nucleus (anteroposterior (AP) −1.35 mm, mediolateral (ML) 0 mm, dorsoventral (DV) −3.6 mm relative to bregma). Bilateral screw electrodes were positioned over the somatosensory (barrel) cortex (AP −1.0 mm, ML ±3.0 mm) for cortical EEG recordings. For sleep-stage determination, EMG activity was recorded from Teflon-coated wire electrodes inserted bilaterally into the nuchal muscles; epidural screw electrodes placed behind the lambda on each side of the midline served as the ground (right) and reference (left) electrodes. All electrodes were connected to a miniature head-mounted connector and secured to the skull with dental cement. Mice received flunixin meglumine (Banamine; 2.5 mg/kg, i.p.) immediately after surgery and every 24h for 48h. After surgery, mice were housed individually in recording chambers under a 14:10 h light–dark cycle with ad libitum access to food and water, and were allowed at least 7 days to recover before recordings began. After recovery and habituation to the recording environment, mice were connected to the recording apparatus and allowed to move freely throughout the recording chamber (Figure 1a). Continuous, synchronized electrophysiological and video recordings were acquired with a Pinnacle recording system (Pinnacle Technology Inc., Lawrence, KS, USA), as described previously^20–22^. Cortical EEG, thalamic local field potential (LFP), and EMG signals were recorded simultaneously in freely behaving mice, amplified, and digitized at 2000 Hz with acquisition filtering between 0.5 and 500 Hz. Cortical EEG was recorded bilaterally from the barrel cortex and thalamic LFP from the CMT, which permitted simultaneous assessment of regional spectral activity and of corticocortical and thalamocortical phase synchronization across wakefulness and NREM sleep.

Upon completion of the experiments, thalamic electrode locations were verified histologically using procedures described previously, visualized with bright-field microscopy, and confirmed against a mouse brain atlas (The Mouse Brain in Stereotaxic Coordinates, Paxinos and Franklin, 4^th^ edition). Animals in which the thalamic electrode was located outside the CMT were excluded from thalamic analyses. The cortical screw placement was also verified.

### Automated Sleep-Wake Scoring

Sleep-wake states were scored with IntelliSleepScorer, a machine-learning-based automated sleep-stage scoring system developed for mouse EEG/EMG recordings^23^. IntelliSleepScorer uses a light gradient boosting machine (LightGBM) classifier and assigns each epoch to wakefulness, NREM sleep, or REM sleep^23^. The published model was trained on 5,776h of EEG/EMG recordings from 124 mice and achieved 95.2% overall classification accuracy and a Cohen’s κ of 0.91 against expert human scoring, using an acquisition configuration comparable to that used here^23^.

Continuous recordings were exported to EDF format and analyzed with the 1-EEG-plus-EMG model. Vigilance states were classified in 10s epochs, the default epoch duration used to develop the model. Wakefulness was characterized by low-amplitude, high-frequency EEG activity with elevated EMG tone; NREM sleep by high-amplitude, low-frequency EEG activity with prominent delta power (0.5-4 Hz) and reduced EMG activity; and REM sleep by low-amplitude, mixed-frequency EEG activity, increased theta power, and near-complete muscle atonia. Automated scoring output was inspected visually by an investigator blinded to group assignment, and epochs containing recording artifacts or uncertain classifications were corrected manually in Sirenia Sleep PRO (version 1.6.1, Pinnacle Technology Inc., Lawrence, KS, USA); on average, less than 5% of epochs were identified as artifact-containing based on the standard deviation of the signal and high amplitude, usually head movement-based.

Sleep architecture parameters - cumulative time in wakefulness, NREM sleep, and REM sleep; episode number; mean episode duration; and state-transition frequency - were computed either in 1h bins across the 24h recording or averaged across the acute, dark phase, and light phase intervals, and were compared between saline- and morphine-treated conditions.

### Spectral Analysis

Spectral power was calculated separately for wakefulness, NREM sleep, and REM sleep from barrel cortex EEG and CMT LFP recordings; only epochs classified as the corresponding vigilance state contributed to each state-specific analysis. Power spectra were computed for artifact-free 10s epochs using fast Fourier transform (FFT)-based methods implemented in Brainstorm^24^ and averaged within each vigilance state. Both absolute power, expressed as power spectral density (PSD, µV²/Hz), and relative power (percentage of total power) were quantified within conventional frequency bands: delta (0.5-4 Hz), theta (4-8 Hz), alpha (8-15 Hz), beta (15-30 Hz), and low gamma (30-50 Hz). For time-resolved analyses, spectral power was averaged within 1-h bins across the 24-h recording.

PSD values were converted to decibels for statistical analysis:

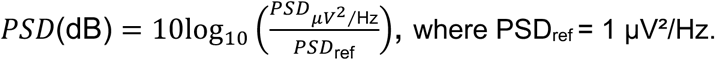

To quantify drug-induced changes in spectral power while controlling for inter-animal variability, we computed within-mouse effects relative to vehicle. For each vigilance state, average PSD values were expressed as ΔdB relative to saline: 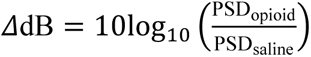 or across time bins:

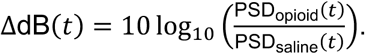

To determine whether morphine altered the relationship between vigilance-state expression and regional oscillatory activity, we correlated the time spent in a state with frequency-specific spectral power in that same state. For each animal, time spent in wakefulness during each hour was correlated with the corresponding wake spectral power, and time spent in NREM sleep with the corresponding NREM spectral power. Correlations were computed separately for the barrel cortex and CMT, and for the delta, theta, alpha, beta, and low-gamma bands, in GraphPad Prism (version 11.2, GraphPad Software, Boston, Massachusetts USA). Pearson correlation coefficients were transformed with Fisher’s r-to-z transformation before comparison between conditions, which stabilizes the variance of the coefficients and permits comparison among the saline, first morphine exposure, fourth morphine exposure, and abstinence conditions:

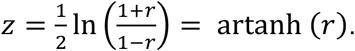

The correlations are computed on hourly values and pooled across animals for overall reporting of Pearson’s r, slopes, and statistical significance. For additional analysis, 24-hourly bins were entered for each correlation, computed per animal, and transformed to Fisher’s z for comparison between conditions.

Functional phase synchronization was assessed using the weighted phase-lag index (wPLI). wPLI quantifies the consistency of non-zero phase-lag relationships between two signals, weighting each phase difference by its magnitude, and is therefore comparatively insensitive to the zero-lag coupling that arises from volume conduction and common-source effects^18^. wPLI was calculated using Hilbert transformation in Brainstorm^24^ separately for wakefulness and NREM sleep, between the left and right cortical electrodes to index corticocortical synchronization and between the CMT and barrel cortex to index thalamocortical synchronization. Values were computed in 10s windows to align with the scoring epochs and were subsequently averaged within each hour of the 24h recording or across the acute, dark phase, and light phase intervals. Because wPLI measures statistical phase synchronization rather than directional information flow, these analyses are interpreted as measures of functional phase coordination rather than of anatomical or causal connectivity. The wPLI was computed for the delta, theta, alpha, beta, and low-gamma frequency bands using the same band definitions as in the spectral analysis, for wake and NREM sleep. Both averaged, and hourly analysis of wPLI were analyzed across the first/fourth injection and abstinence.

### Statistical Analysis

Statistical analyses were performed in GraphPad Prism (version 11.2, GraphPad Software, Boston, Massachusetts USA). Vigilance-state occupancy, behavioral measures, spectral power, and wPLI were analyzed using a two-way repeated-measures (RM) ANOVA or a mixed-effects model when missing values were present, with treatment or exposure condition and time or vigilance state as factors, followed by Šídák multiple-comparison tests where appropriate. Multiple comparisons were analyzed separately for each of the five frequency bands across three vigilance states in the cortex and thalamus. For correlation analyses, Pearson correlation coefficients were calculated for each frequency band and vigilance state and then transformed to Fisher z values before between-condition comparisons. Data are presented as mean±SEM. Statistical significance was defined as p<0.05, and significance levels in figures are indicated as *p<0.05, **p<0.01, ***p<0.001, and ****p<0.0001. Both male and female mice were included, and since no sex difference in the morphine effect was observed, their data were combined for analysis. One animal was excluded due to poor placement of the deep electrode. Additionally, one cortical recording was lost, and wPLI analysis was performed on data from 6 animals.

## RESULTS

Statistical test results and sample sizes are given in the figure legends.

### Sleep-wake architecture across time after repeated morphine administration

EEG/LFP/EMG recordings were obtained across the 24h following the first/fourth injection, and the abstinence day (Figure 1a). Animals received saline at ZT9 on four consecutive days, and after the recovery period, received morphine at ZT9 on four consecutive days. Vigilance states were quantified across the acute effect (5h after injection, during light phase), the dark phase, and the following light phase.

After the first morphine injection, wakefulness increased during the post-injection acute phase (ZT11-13), and NREM sleep was correspondingly suppressed (Figure 1b,c). REM sleep was absent during the post-injection period, with a slight rebound at ZT0 (Figure 1d). During the early dark phase, wakefulness declined and NREM sleep rose in the morphine group (ZT16, Figure 1b,c). Interestingly, in the saline group, wakefulness decreased later in the dark phase (ZT21; Figure 1b,c). During the light phase distributions were comparable across groups, indicating that the wake-promoting effect was transient. After the fourth injection, wakefulness was again greater immediately after morphine injection, with a corresponding reduction in NREM sleep (ZT11-13; Figure 1e,f), but the temporal profile differed from the first exposure. During the early dark phase, morphine reduced wakefulness and increased NREM sleep (ZT16,17,19; Figure 1f). REM sleep was absent immediately after injection (ZT12,13) and increased later in the dark phase (ZT22; Figure 1g). During abstinence vigilance-state occupancy did not differ between saline- and morphine-withdrawn animals in either phase (Figure 1h,i,j).

### Behavior and sleep-wake architecture during the acute morphine effect

Morphine-treated animals were more active than controls on all four testing days, with increased distance traveled (Figure 2a), decreased immobile time (Figure 2b), and higher mean speed (Figure 2c) over the 1h observation period. Trajectory plots and heat maps showed broader arena exploration in the saline group, with thigmotaxic behavior in morphine-treated animals (Figure 2d,e). During abstinence, paw withdrawal thresholds were reduced about 20% in the morphine group relative to baseline and saline controls (Figure 2f), consistent with opioid-induced hyperalgesia. Averaged sleep architecture was also altered during the acute phase (Figure 2g,h,i). The number of wake, NREM, and REM episodes decreased after the first injection (Figure 2g left); mean wake episode duration increased after both injections (Figure 2h left, middle). Cumulative wake time increased, while NREM time decreased after injections (Figure 2i left, middle). Fewer and longer wake episodes indicate consolidation of wakefulness.

**Figure 2.**
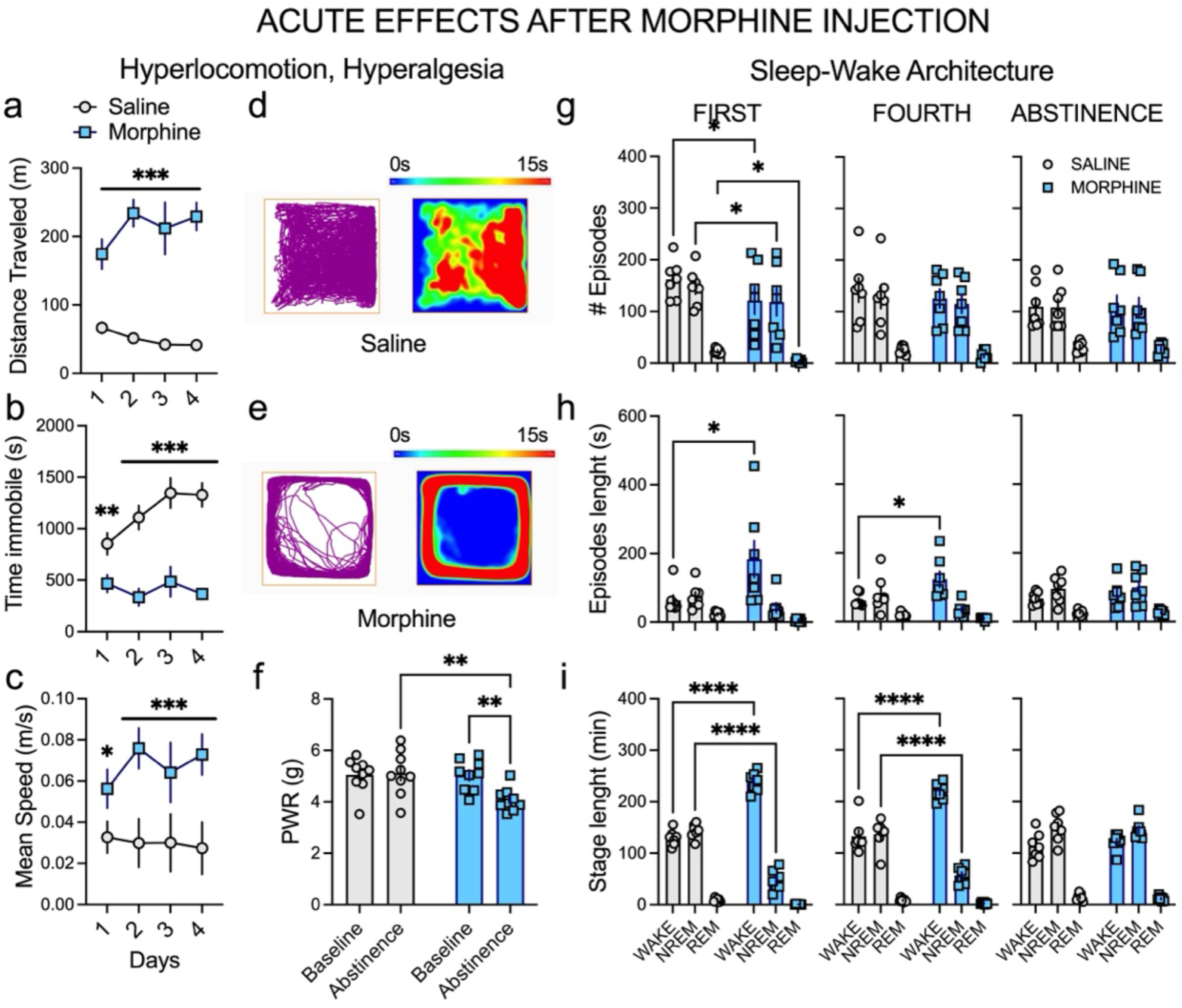
During the light phase after the injections, morphine induces hyperlocomotion, alters sleep-wake architecture, and induces hyperalgesia during abstinence. (a) Total distance travelled during four consecutive testing days in saline- and morphine-treated animals under acute light conditions. Morphine-treated animals exhibited significantly greater locomotor activity than saline controls; two-way RM ANOVA: Time F_3,21_=1.02, p=0.04; Morphine F_1,7_=56.29, p=0.0001, Interaction F_3,21_=3.06, p=0.05, Šídák’s post hoc presented in the figure. (b) Total immobility time across the four testing days. Morphine administration significantly reduced immobility compared with saline-treated animals; two-way RM ANOVA: Time F_3,21_=5.00, p=0.009; Morphine F_1,7_=37.25, p=0.0005, Interaction F_3,21_=3.74, p=0.03, Šídák’s post hoc presented in the figure. (c) Mean locomotor speed across four testing days, showing elevated speed in morphine-treated animals relative to saline controls; two-way RM ANOVA: Time F_3,21_=1.04, p=0.395; Morphine F_1,7_=3.66, p=0.097, Interaction F_3,21_=1.60, p=0.219, Šídák’s post hoc presented in the figure. (d) Representative locomotor trajectory (left) and occupancy heat map (right) of a saline-treated animal. Warmer colors indicate longer dwell times. (e) Representative locomotor trajectory and occupancy heat map of a morphine-treated animal, demonstrating increased peripheral exploration and reduced stationary behavior. (f) Mechanical paw abstinence response (PWR) was measured at baseline and during abstinence. Morphine-treated animals exhibited reduced mechanical withdrawal thresholds during abstinence, consistent with hyperalgesia, whereas saline controls remained stable; two-way RM ANOVA: Abstinence F_1,16_=4.01, p=0.06; Morphine F_1,16_=5.73, p=0.03, Interaction F_1,16_=5.35, p=0.03, Šídák’s post hoc presented in the figure. (g-i) Quantification of sleep-wake episodes during the acute effect after the first (left), fourth injection (middle), and abstinence (right). Bars represent mean values with individual data points overlaid for saline (circles) and morphine (squares). (g) Mean number of episodes. (h) Mean episode duration for vigilance states (i) Total time spent in wakefulness, NREM and REM sleep. Two-way RN ANOVA: number of episodes after the <u>first injection</u>: Stage F_2,12_=53.08, p<0.0001; Morphine F_1,6_=2.83, p=0.143; Interaction F_2,12_=0.54, p=0.597; episodes length after the <u>first injection</u>: Stage F_2,12_=11.17, p=0.002; Morphine F_1,6_=1.32, p=0.295; Interaction F_2,12_=6.29. p=0.013; episodes length after the <u>fourth injection</u>: Stage F_2,12_=24.66, p<0.0001; Morphine F_1,6_=0.008, p=0.931; Interaction F_2,12_=9.22 p=0.004; stage length after the first injection: Stage F_2,12_=324.7, p<0.0001; Morphine F_1,6_=400.0, p<0.0001; Interaction F_2,12_=154.6 p<0.0001; stage length after the <u>fourth injection</u>: Stage F_2,12_=132.2, p<0.0001; Morphine F_1,6_=0.033, p=0.861; Interaction F_2,12_=56.38 p<0.0001; Šídák’s post hoc is presented in the figure. Data are presented as mean ± SEM, N=8 for locomotion testing, N=9 for PWR, N=7 animals for sleep studies

### Sleep-wake architecture during the dark and light phases

Sleep architecture was also evaluated separately for the dark and light phases. During the dark phase, wake, and NREM episode numbers increased after the fourth injection (Figure S1a, middle), and episode duration decreased after both injections (Figure S1b, left and middle), indicating reduced sleep continuity. Cumulative time was redistributed after the fourth injection toward less wakefulness and more NREM sleep (Figure S1c, middle). These differences were absent during abstinence.

During the light phase, wake, and NREM s the fourth injection (Figure S1d middle), and the wake difference persisted into abstinence (Figure S1d right). NREM episode duration increased after the first injection (Figure S1e left), which also decreased cumulative wakefulness and increased cumulative NREM sleep (Figure S1f left). Normalization was therefore partial: hourly occupancy recovered, while the number of light phase wake episodes remained elevated.

### Cortical oscillatory activity during wakefulness and NREM sleep

Representative spectrograms showed distinct cortical changes after the first injection (acute effect; Figure 3a). Morphine redistributed cortical spectral power across bands during the acute interval, and both subsequent phases (Figure 3b). Table 1 gives the direction of change for every band, state, and exposure. Three patterns organize the acute effects. Delta and low-gamma power increased during both wakefulness and NREM sleep after each injection, and the delta increase persisted into abstinence. Alpha power moved in opposite directions with exposure history, decreasing after the first injection and increasing after the fourth during wakefulness. Theta and beta power changed mainly with repeated exposure and abstinence. REM effects were the smallest, restricted to increased delta, beta, and low-gamma power after the fourth injection (Figure 3b top, Table 1). Across the dark and light phases, cortical power was elevated in most bands, more so after repeated injections (Figure 3b middle, bottom; Tables S1, S2).

**Figure 3.**
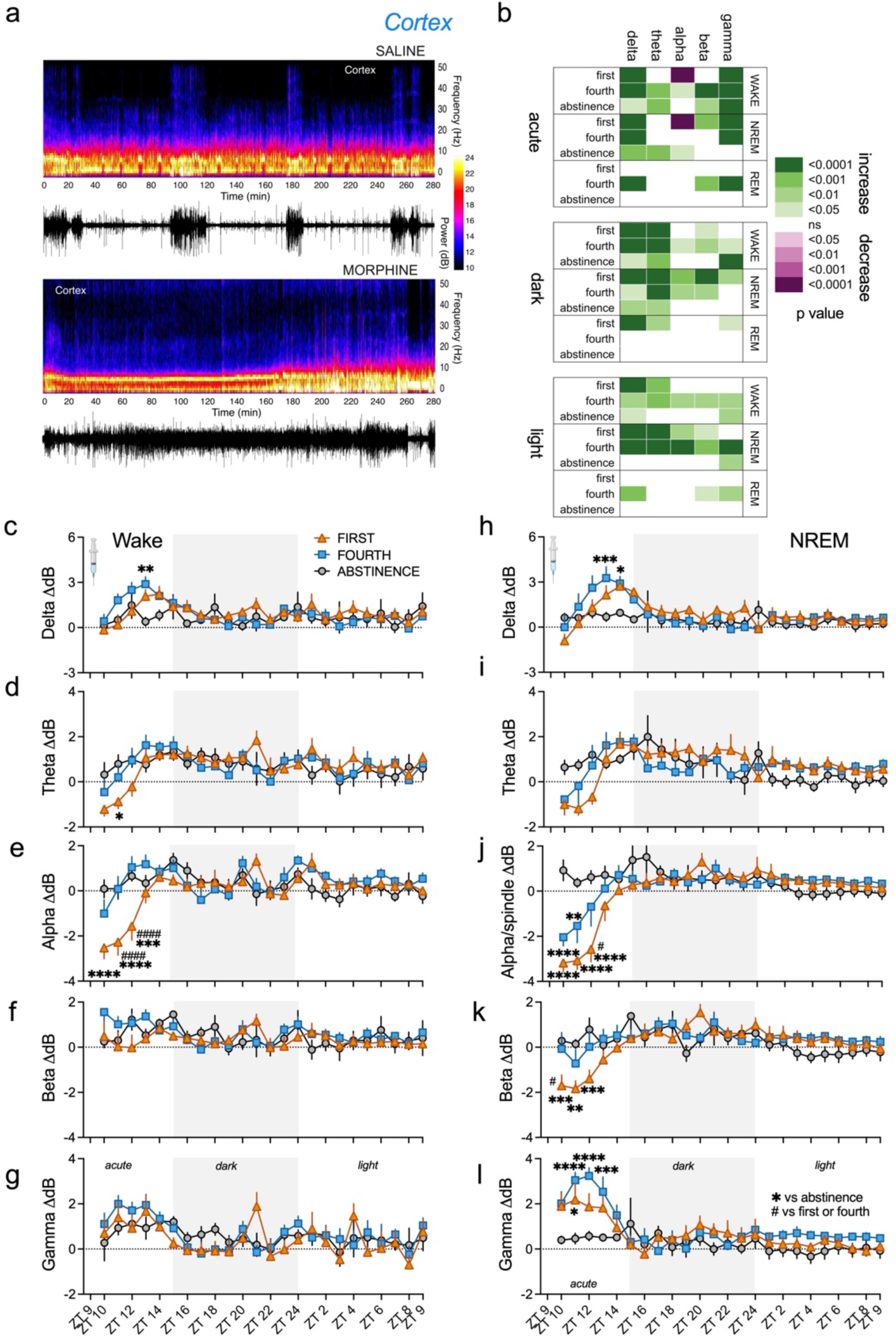
Morphine alters cortical oscillatory activity during wakefulness, NREM sleep, and REM sleep after injections. (a) Representative time-frequency spectrograms showing cortical (EEG) activity following the first administration of saline (top) or morphine (bottom). The lower traces show representative EMG recordings. Color scale indicates spectral power (dB). (b) Summarized statistical analysis of averaged cortical wake/NREM/REM spectral power during the acute effect, dark, and light phases (top, middle, bottom, respectively). Cortical spectral power was analyzed across the delta, theta, alpha, beta, and low gamma frequency bands. (c-l) Normalized EEG spectral power changes during wakefulness (c-g) and NREM (h-l) across barrel cortex. Changes in normalized power (ΔdB) are shown across recording time points (ZT) for the cortex during wake/NREM sleep in the delta (c, h), theta (d, i), alpha (e, j), beta (f, k), and gamma (g, l) frequency bands. Data are presented for the first exposure (orange triangles), the fourth injection (blue squares), and during abstinence (gray circles). Symbols represent group means and SEM. The horizontal dotted line at 0 ΔdB denotes no change relative to the saline. Gray-shaded regions indicate the dark phase, and syringe symbols indicate the timing of experimental saline/morphine administration. The statistically significant Šídák post hoc result was presented as *compared to abstinence or ^#^compared to first or fourth injection. Two-way RM ANOVA results: <u>Delta wake</u>: Time F_23,138_=4.18, p<0.0001; Condition F_2,12_=0.40, p=0.678; Interaction F_46,276_=1.38, p=0.006; <u>Theta wake</u>: Time F_23,138_=5.61, p<0.0001; Condition F_2,12_=0.005, p=0.995; Interaction F_46,276_=2.17, p=0.01; <u>Alpha wake</u>: Time F_23,138_=6.81, p<0.0001; Condition F_2,12_=2.80, p=0.213; Interaction F_46,276_=3.00, p<0.0001. Mixed-effects model (REML) results: <u>Delta NREM</u>: Time F_23,138_=8.14, p<0.0001; Condition F_2,12_=1.76, p=0.213; Interaction F_46,276_=1.38, p<0.0001; <u>Alpha NREM</u>: Time F_23,138_=12.72, p<0.0001; Condition F_2,12_=2.30, p=0.143; Interaction F_46,276_=5.39, p<0.0001; <u>Beta NREM</u>: Time F_23,138_=5.72, p<0.0001; Condition F_2,12_=0.68, p=0.523; Interaction F_46,276_=2.61, p<0.0001; <u>Gamma NREM</u>: Time F_23,138_=8.77, p<0.0001; Condition F_2,12_=5.09, p=0.025; Interaction F_46,275_=2.45, p<0.0001. N=7 animals, *p<0.05, **p<0.01, ***p<0.001, ****p<0.0001; ^#^p<0.05, ^##^p<0.01, ^###^p<0.001, ^####^p<0.0001

**Table 1.**
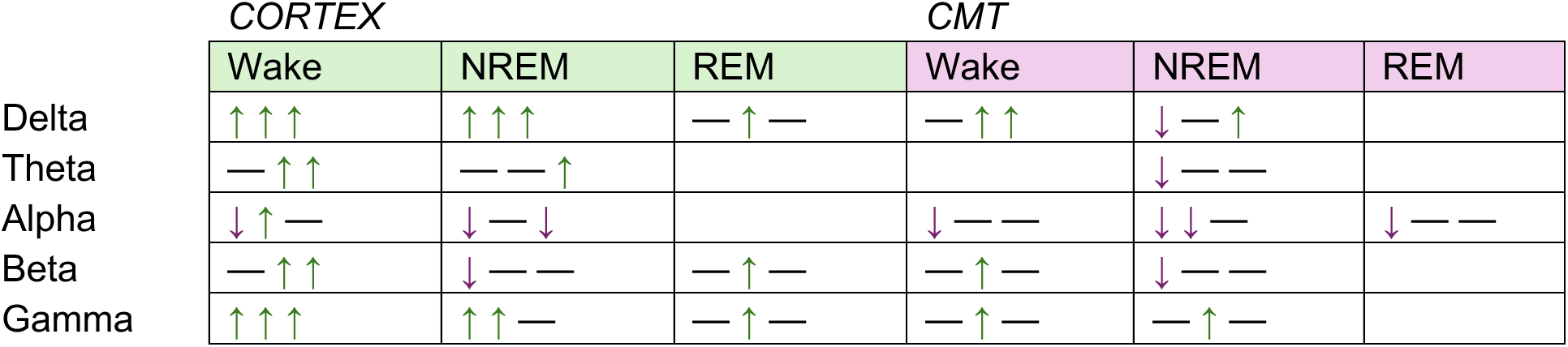
Direction of morphine-induced change in cortical and central medial thalamic (CMT) spectral power during the acute post-injection interval (5 h), by frequency band, vigilance state, and recording site. Each cell gives the first injection / fourth injection / abstinence, in that order. ↑ statistically significant increase, ↓ statistically significant decrease, — no significant change reported. Compiled from Figures 3b and 4b (top row) and Figure S2.

Over the 24h period, the largest changes occurred during the acute interval and the first hours of abstinence (Figure 3c-l). Cortical delta power rose rapidly, and earlier after the fourth than the first injection, in both states (Figure 3c,h). Wake theta power decreased early after the first injection (Figure 3d). Alpha power decreased markedly after the first injection in both states and changed only modestly after the fourth (Figure 3e,j). Beta power showed a transient NREM decrease after the first exposure (Figure 3k). Gamma power was elevated early in all conditions before declining toward baseline during wakefulness (Figure 3g), whereas during NREM sleep repeated injection produced a substantial rise 2-4 h after injection (Figure 3l). Hourly power was otherwise comparable across conditions during the dark and light phases.

### CMT oscillatory activity during wakefulness and NREM sleep

Representative spectrograms showed distinct thalamic changes after the first injection (acute effect; Figure 4a, which also shows the deep electrode confirmation). Morphine redistributed CMT spectral power most clearly after the injections, predominantly during wakefulness and NREM sleep (Figure 4b); directions for every band, state, and condition are given in Table 1. The thalamic pattern differed from the cortical one in two respects. First, the first injection produced predominantly reductions in delta, theta, alpha, and beta power during NREM sleep, and alpha power during wakefulness and REM sleep, whereas cortical delta and low-gamma power increased. Second, repeated exposure shifted the thalamic response toward increases, most consistently in low gamma. During abstinence, only delta power remained elevated in both wakefulness and NREM sleep (Figure 4b top, Table 1). Across the dark and light phases, thalamic power was elevated in most bands except low gamma, more so after repeated injections (Figure 4b middle and bottom; Tables S1, S2).

**Figure 4.**
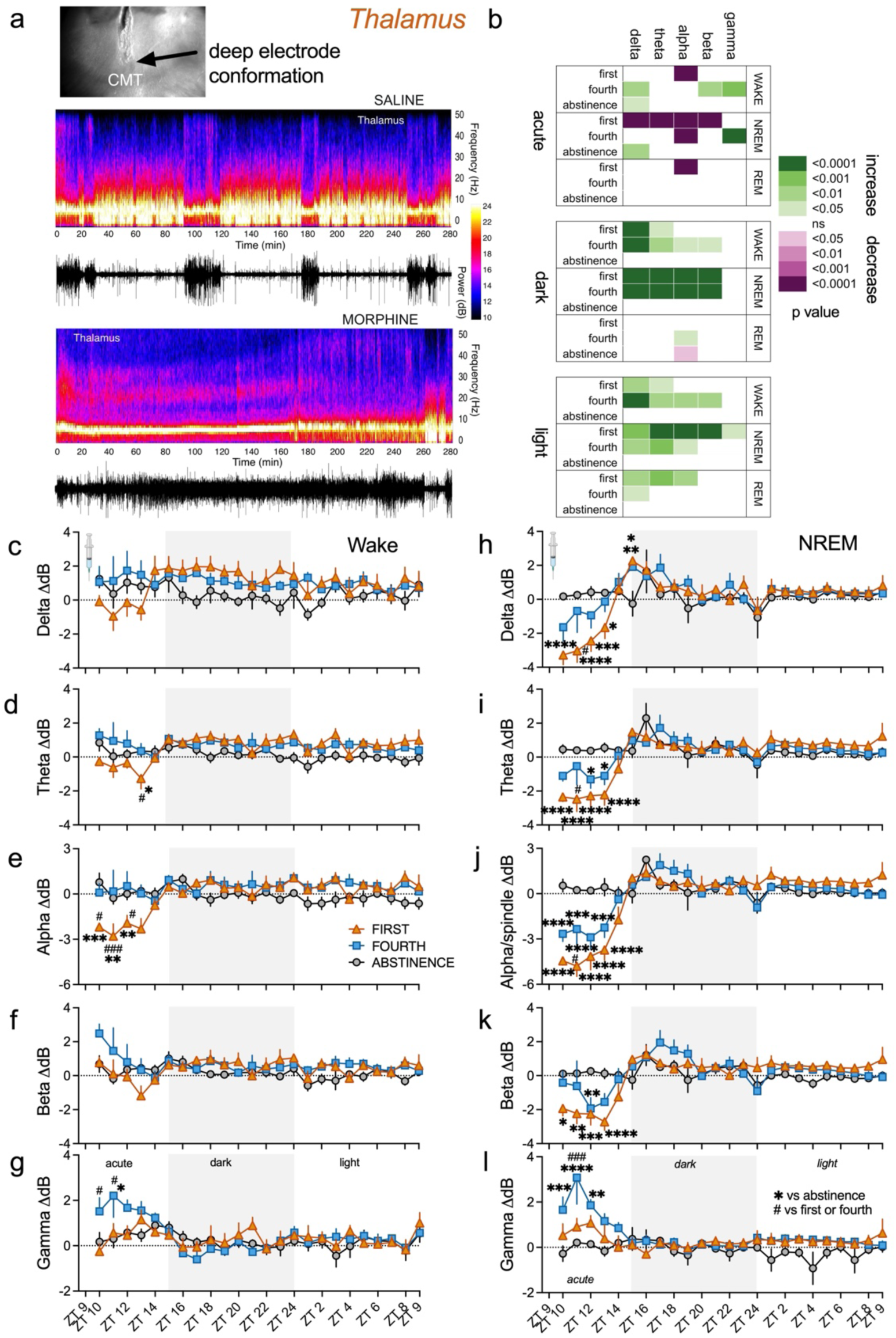
Morphine alters CMT oscillatory activity during wakefulness, NREM sleep, and REM sleep after injections. (a) Representative time-frequency spectrograms showing thalamic (LFP) activity following the first administration of saline (top) or morphine (bottom) with deep electrode conformation, the black arrow shows the end of deep thalamic electrode placement right above the CMT region. The lower traces show representative EMG recordings. Color scale indicates spectral power (dB). (b) Summarized statistical analysis of averaged thalamic wake/NREM/REM spectral power during the acute effect, dark, and light phases (top, middle, bottom, respectively). CMT spectral power was analyzed across the delta, theta, alpha, beta, and low gamma frequency bands. (c-l) Normalized EEG spectral power changes during wakefulness (c-g) and NREM (h-l) across CMT. Changes in normalized power (ΔdB) are shown across recording time points (ZT) for the thalamus during wake/NREM sleep in the delta (c, h), theta (d, i), alpha (e, j), beta (f, k), and low gamma (g, l) frequency bands. Data are presented for the first exposure (orange triangles), the fourth injection (blue squares), and during abstinence (gray circles). Symbols represent group means and SEM. The horizontal dotted line at 0 ΔdB denotes no change relative to the saline. Gray-shaded regions indicate the dark phase, and syringe symbols indicate the timing of experimental saline/morphine administration. The statistically significant Šídák post hoc result was presented as *compared to abstinence or ^#^compared to first or fourth injection. Two-way RM ANOVA results: <u>Theta wake</u>: Time F_23,138_=1.58, p=0.06; Condition F_2,12_=6.32, p=0.013; Interaction F_46,276_=1.95, p=0.0006; <u>Alpha wake</u>: Time F_23,138_=2.39, p=0.001; Condition F_2,12_=3.42, p=0.07; Interaction F_46,276_=2.73, p<0.0001. Mixed-effects model (REML) results: <u>Gamma wake</u>: Time F_23,138_=4.00, p<0.0001; Condition F_2,12_=0.67, p=0.529; Interaction F_46,252_=1.58, p=0.015; <u>Delta NREM</u>: Time F_23,138_=5.51, p<0.0001; Condition F_2,12_=1.20, p=0.336; Interaction F_46,235_=12.91, p<0.0001; <u>Theta NREM</u>: Time F_23,138_=7.15, p<0.0001; Condition F_2,12_=20.80, p=0.472; Interaction F_46,256_=3.80, p<0.0001; <u>Alpha NREM</u>: Time F_23,138_=11.89, p<0.0001; Condition F_2,12_=7.27, p=0.009; Interaction F_46,256_=6.24, p<0.0001; <u>Beta NREM</u>: Time F_23,138_=5.52, p<0.0001; Condition F_2,12_=0.67, p=0.531; Interaction F_46,256_=3.37, p<0.0001; <u>Gamma NREM</u>: Time F_23,138_=3.94, p<0.0001; Condition F_2,12_=7.98, p=0.006; Interaction F_46,235_=2.20, p<0.0001. N=7, *p<0.05, **p<0.01, ***p<0.001, ****p<0.0001; ^#^p<0.05, ^##^p<0.01, ^###^p<0.001, ^####^p<0.0001

The largest CMT changes also occurred during the acute interval and the first hours of abstinence. During wakefulness, thalamic delta power did not differ between conditions, although delta power was modestly elevated after the fourth injection (Figure 4c). Theta power showed a brief early reduction after the first injection (Figure 4d), and alpha power showed marked early suppression, reaching approximately −3 ΔdB over the first 3h (Figure 4e). Beta power was initially highest following the repeated injections, with no differences between conditions (Figure 4f). Gamma power was initially elevated after the fourth injection relative to the first and to abstinence (Figure 4g). During NREM sleep, delta power decreased early after the first injection and modestly after the fourth, then increased during the first hours of the dark phase (Figure 4h). Theta power was reduced after the first injection and modestly after the fourth (Figure 4i). Alpha power again showed marked early suppression, approximately −4 to −5 ΔdB after the first injection and −3 ΔdB after the fourth (Figure 4j). Beta power decreased early after both injections (Figure 4k). Gamma power was initially elevated, most clearly after the fourth injection (Figure 4l). The first exposure, therefore, reduced thalamic delta and alpha power during NREM sleep, whereas repeated exposure primarily elevated CMT gamma power, as in the cortex.

### Correlations between spectral power and time spent in wakefulness or NREM sleep

Wake or NREM duration (% time) was correlated with absolute power (PSD) for each condition across the 24h period in 1h bins (plots per band, Figures S2–S7; summary in Figure 5a). Table 2 shows the slope comparisons, and Table 3 shows the morphine-saline differences in Fisher’s z-transformed coefficients. Associations were strongest at delta, theta, and alpha, and morphine altered both their direction and magnitude differently after the first injection, the fourth, and abstinence (Figure 5a). Under saline conditions, cortical power correlated positively with wake duration, particularly in the theta, alpha, and beta bands; after morphine, negative associations were observed in the same bands, with a different pattern after repeated exposure (Figure 5a, top). Thalamic correlations showed negative correlations in wake-delta and wake-theta under saline, whereas gamma thalamic activity was more consistently positive. Morphine altered these delta/theta correlations following repeated injections and abstinence (Figure 5a top). NREM correlations were likewise treatment- and exposure-dependent, with cortical and thalamic relationships differing in direction, with the cortical ones being more negative (Figure 5a bottom). Interestingly, the duration of NREM sleep was negatively correlated with cortical delta power after the fourth saline injection and morphine-related abstinence. In CMT, positive NREM-delta correlation was seen just in morphine-treated animals after both injections. Additionally, cortical gamma power showed a negative correlation with NREM duration, in the cortex and thalamus, primarily under morphine (Figure 5a bottom).

**Figure 5.**
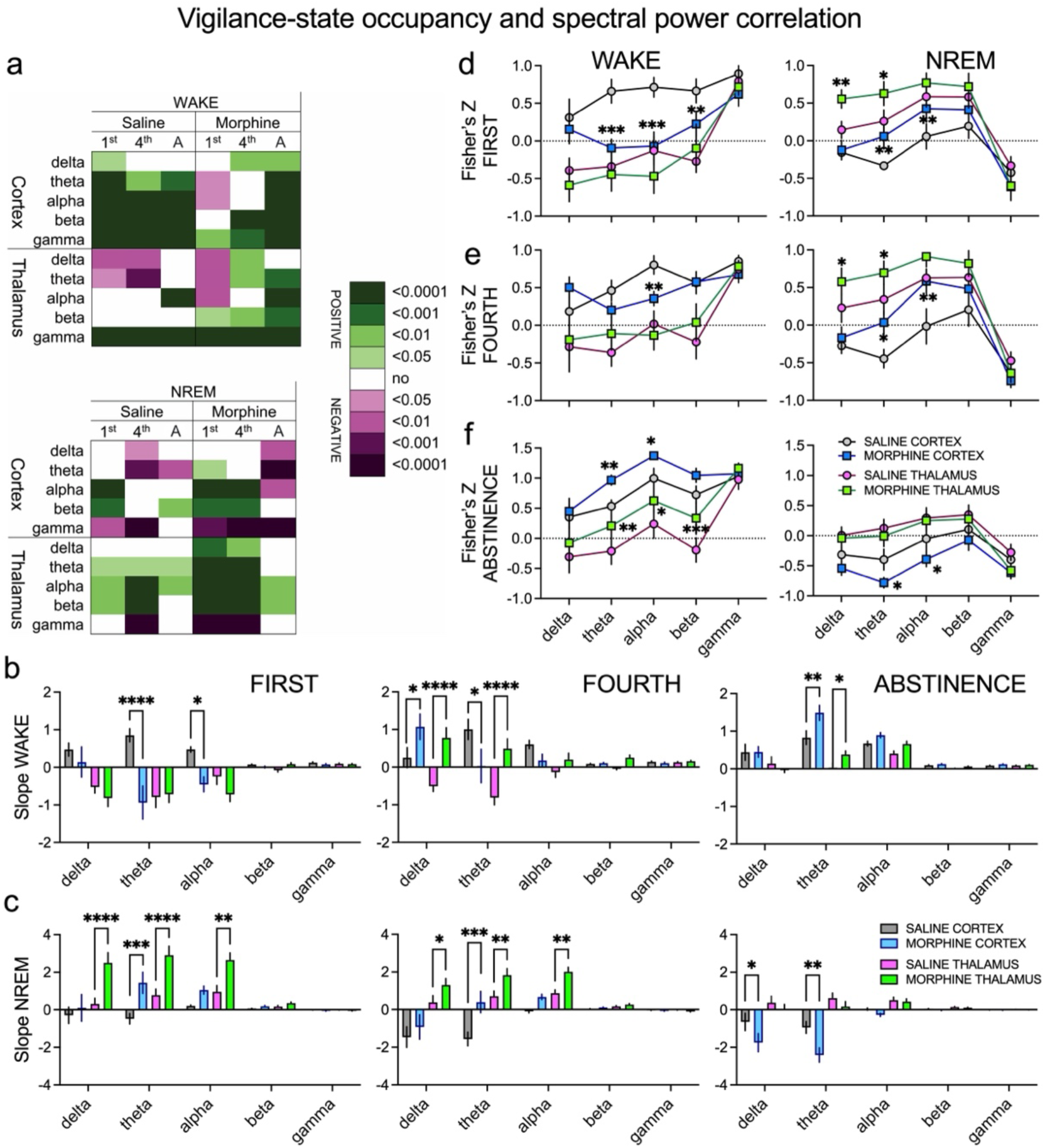
Correlations between cortical and thalamic oscillatory power and vigilance states during repeated morphine treatment and abstinence. (a) Summary heat maps showing statistically significant Pearson’s r correlation coefficients between spectral power in different frequency bands (delta, theta, alpha, beta, and gamma) and the percentage of time spent in wakefulness (wake) or NREM sleep. Correlations were calculated separately for cortical and thalamic recordings after the first (1^st^), fourth injection (4^th^), and abstinence (A) in saline- and morphine-treated animals. Green colors indicate significant positive correlations, whereas purple colors indicate significant negative correlations. (b) Slope analysis during wakefulness after the first (left), fourth injection (middle), and abstinence (right) in cortex, saline (gray) and morphine (blue), and thalamus, saline (pink) and morphine (green). Two-way ANOVA for <u>first injection Cortex</u>: Frequency F_4,60_=0.785, p=0.540, Morphine F_1,60_=21.58, p<0.0001, Interaction F_4,60_=5.89, p=0.0005; <u>fourth injection Thalamus</u>: Frequency F_4,60_=1.20, p=0.319, Morphine F_1,60_=40.39, p<0.0001, Interaction F_4,60_=6.94, p=0.0001; <u>fourth Cortex:</u> Frequency F_4,60_=2.43, p=0.057, Morphine F_1,60_=0.71, p=0.40, Interaction F_4,60_=4.35, p=0.004; <u>abstinence Thalamus:</u> Frequency F_4,60_=11.06, p<0.0001, Morphine F_1,60_=3.29, p=0.07, Interaction F_4,60_=2.80, p=0.034, <u>abstinence Cortex:</u> Frequency F_4,60_=27.43, p<0.0001, Morphine F_1,60_=6.10, p=0.016, Interaction F_4,60_=2.54, p=0.049; Šídák’s post hoc was presented in the figures. (c) Slope analysis during NREM sleep after the first (left), fourth injection (middle) and abstinence (right) in cortex, saline (gray) and morphine (blue), and thalamus, saline (pink) and morphine (green). Two-way ANOVA for <u>first injection Thalamus</u>: Frequency F_4,60_=15.67, p<0.0001, Morphine F_1,60_=37.86, p<0.0001, Interaction F_4,60_=5.89, p=0.0005; <u>first injection Cortex</u>: Frequency F_4,60_=1.79, p=0.142, Morphine F_1,60_=8.99, p=0.004, Interaction F_4,60_=2.67, p=0.041; <u>fourth injection Thalamus</u>: Frequency F_4,60_=15.24, p<0.0001, Morphine F_1,60_=18.24, p<0.0001, Interaction F_4,60_=3.08, p=0.022; <u>fourth injection Cortex</u>: Frequency F_4,60_=5.85, p=0.0005, Morphine F_1,60_=9.10, p=0.004, Interaction F_4,60_=2.62, p=0.045; <u>abstinence Cortex:</u> Frequency F_4,60_=17.19, p<0.0001, Morphine F_1,60_=12.43, p=0.0008, Interaction F_4,60_=2.98, p=0.026; Šídák’s post hoc was presented in the figures. (d) Fisher’s z analysis for wake (left) and NREM (right) in the cortex and thalamus after the first injection. Two-way RM ANOVA for <u>wake episodes Cortex</u>: Frequency F_4,24_=5.08, p=0.004, Morphine F_1,6_=13.75, p=0.01, Interaction F_4,24_=5.23, p=0.003; <u>NREM Cortex:</u> Frequency F_4,24_=23.04, p<0.0001, Morphine F_1,6_=1.29, p=0.300, Interaction F_4,24_=5.71, p=0.002; <u>NREM Thalamus:</u> Frequency F_4,24_=21.12, p<0.0001, Morphine F_1,6_=4.84, p=0.07, Interaction F_4,24_=5.58, p=0.002; Šídák’s post hoc was presented in the figures. (e) Fisher’s z analysis for wake (left) and NREM (right) in the cortex and thalamus after the fourth injection. Two-way RM ANOVA for <u>wake episodes Cortex</u>: Frequency F_4,24_=6.00, p=0.002, Morphine F_1,6_=0.75, p=0.42, Interaction F_4,24_=6.06, p=0.002; <u>NREM Cortex:</u> Frequency F_4,24_=22.21, p<0.0001, Morphine F_1,6_=6.21, p=0.05, Interaction F_4,24_=3.37, p=0.025; <u>NREM Thalamus:</u> Frequency F_4,24_=28.15, p<0.0001, Morphine F_1,6_=2.98, p=0.135, Interaction F_4,24_=3.72, p=0.017; Šídák’s post hoc was presented in the figures. (f) Fisher’s z analysis for wake (left) and NREM (right) in the cortex and thalamus during abstinence. Two-way RM ANOVA for <u>wake episodes Cortex</u>: Frequency F_4,24_=12.48, p<0.0001, Morphine F_1,6_=8.58, p=0.026, Interaction F_4,24_=1.92, p=0.140; <u>wake Thalamus:</u> Frequency F_4,24_=10.78, p<0.0001, Morphine F_1,6_=10.32, p=0.018, Interaction F_4,24_=1.46, p=0.244; <u>NREM Cortex:</u> Frequency F_4,24_=11.74, p<0.0001, Morphine F_1,6_=10.21, p=0.019, Interaction F_4,24_=0.57, p=0.683; Šídák’s post hoc was presented in the figures. N=7 animals

**Table 2.**
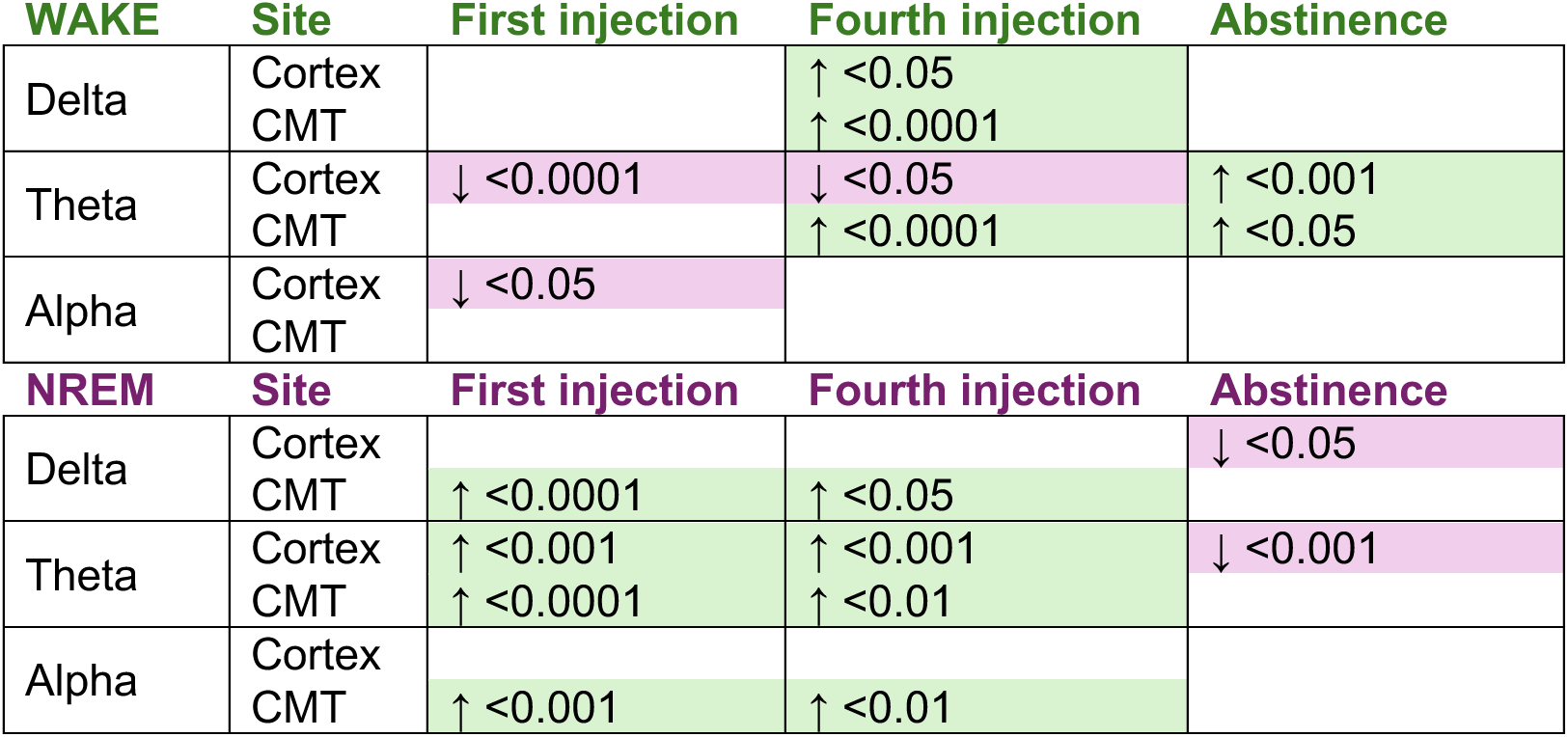
Morphine-induced change in the slope of the relationship between vigilance-state occupancy and band-specific spectral power (Figure 5b-c). Each entry compares morphine with saline within the same recording site; arrows give the direction of the morphine slope relative to saline, followed by the significance level. CMT, central medial thalamus.

**Table 3.**
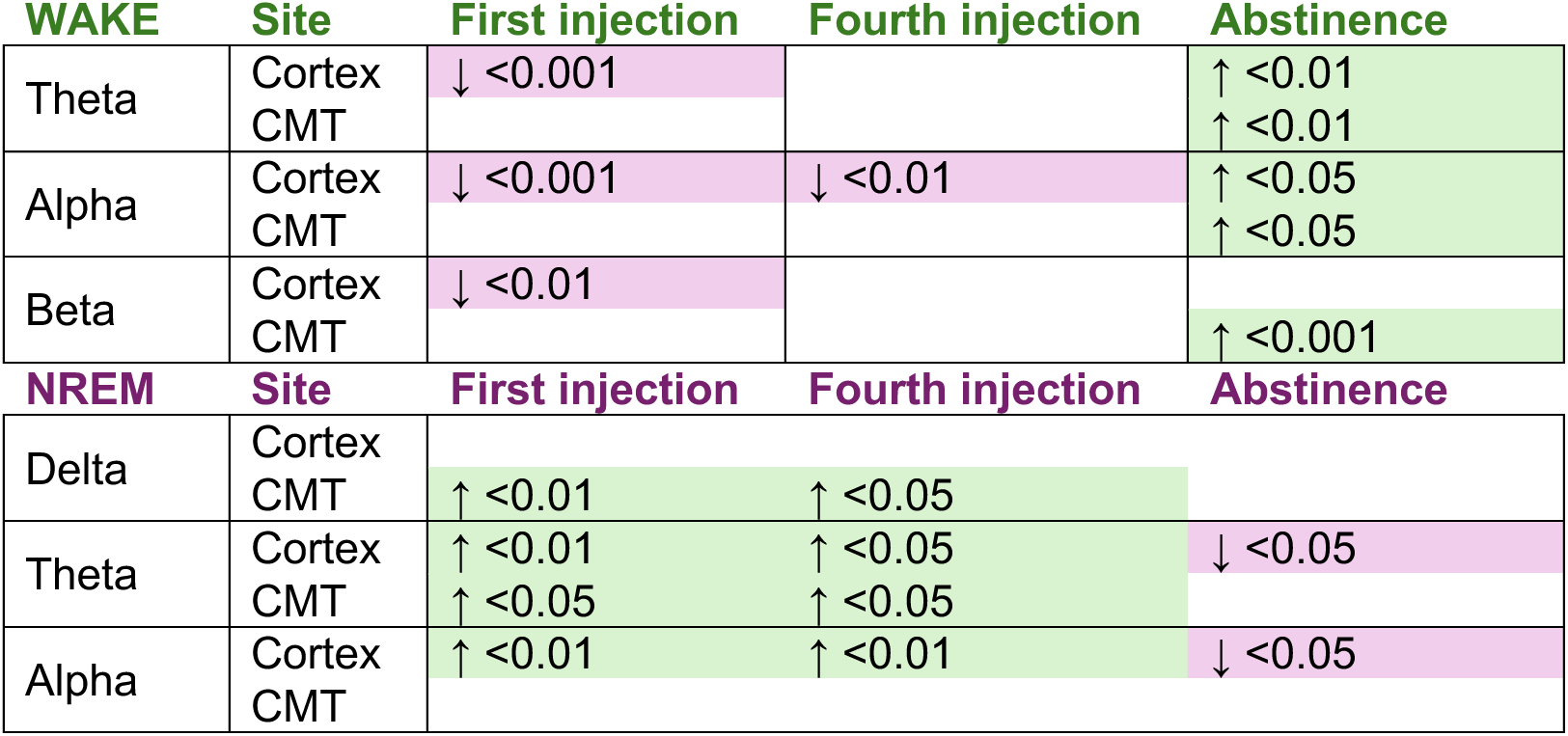
Morphine-induced change in the correlation between vigilance-state occupancy and band-specific spectral power, expressed as the difference in Fisher’s z-transformed correlation coefficients between morphine- and saline-treated mice (Figure 5d-f). Arrows give the direction of the morphine value relative to saline (↑ more positive, ↓ more negative), followed by the significance level. The site indicates whether the comparison was made within the cortical or within the central medial thalamic (CMT). All entries were read directly from the significance annotations in Figure 5d-f.

Slope analyses showed treatment-dependent differences in both states (Figure 5b,c; Table 2). For wake duration, effects after the first injection were concentrated in theta and alpha, the cortical theta slope shifting from positive under saline to negative after morphine (Figure 5b left). After the fourth injection, delta slopes increased in both regions while theta slopes dissociated, increasing in the thalamus and decreasing in the cortex (Figure 5b middle). During abstinence, the main effect was in theta, more positive in the cortex after the morphine, and increased in CMT (Figure 5b right). For NREM duration, both injections produced comparable differences at delta, theta, and alpha: the thalamic delta slope was markedly more positive, theta slopes increased in both regions, the thalamic alpha slope increased, while cortical delta and alpha slopes were unchanged (Figure 5c left, middle). During abstinence, the cortical delta and theta slopes decreased compared to saline, showing negative values. (Figure 5c right).

Fisher’s z analyses supported these frequency-specific effects (Figure 5d-f; Table 3). After the first injection, morphine decreased cortical theta, alpha, and beta correlations during wakefulness, with no thalamic comparison reaching significance (Figure 5d left). During NREM sleep, it increased CMT delta and theta and cortical theta and alpha correlations (Figure 5d right). After the fourth injection, the wake reduction was confined to cortical alpha (Figure 5e left), and the NREM pattern resembled that after the first injection (Figure 5e) right. During abstinence, the direction reversed: wake theta and alpha correlations increased in both regions, together with thalamic beta (Figure 5f left), whereas cortical theta and alpha correlations decreased during NREM sleep (Figure 5f right).

### Corticocortical synchronization

Corticocortical synchronization showed frequency- and state-dependent responses after both injections and during abstinence (Figure 6a-j; hourly analyses, wPLI differences from saline were shown and analyzed). The largest changes occurred during the acute and dark phase periods, whereas most bands approached baseline during the light phase. During wakefulness (Figure 6a-e), delta synchronization remained close to baseline (Figure 6a). Theta wPLI rose immediately after injection, then fell below baseline and remained predominantly negative, decreasing during the dark phase after the first injection (Figure 6b). Alpha synchronization decreased early after the first injection and stayed reduced at several dark phase time points (Figure 6c). Beta synchronization rose acutely then declined toward or below baseline, recovering faster after the fourth injection (Figure 6d). Gamma changes were small, rising after injections and remaining elevated throughout the dark phase following the fourth injection (Figure 6e). During NREM sleep (Figure 6f-j), delta synchronization was more variable than during wakefulness: it increased through the dark phase after the first injection, whereas after the fourth, a transient increase was followed by a decrease at the end of the dark phase (Figure 6f). Theta synchronization rose acutely and then recovered toward baseline (Figure 6g), and alpha synchronization was predominantly reduced during the dark phase after the first injection, with no differences between conditions (Figure 6h). Beta synchronization rose acutely, particularly after the first injection, then fell below baseline for much of the dark phase (Figure 6i). Gamma changes were limited, with an acute increase after the fourth injection relative to the first (Figure 6j).

**Figure 6.**
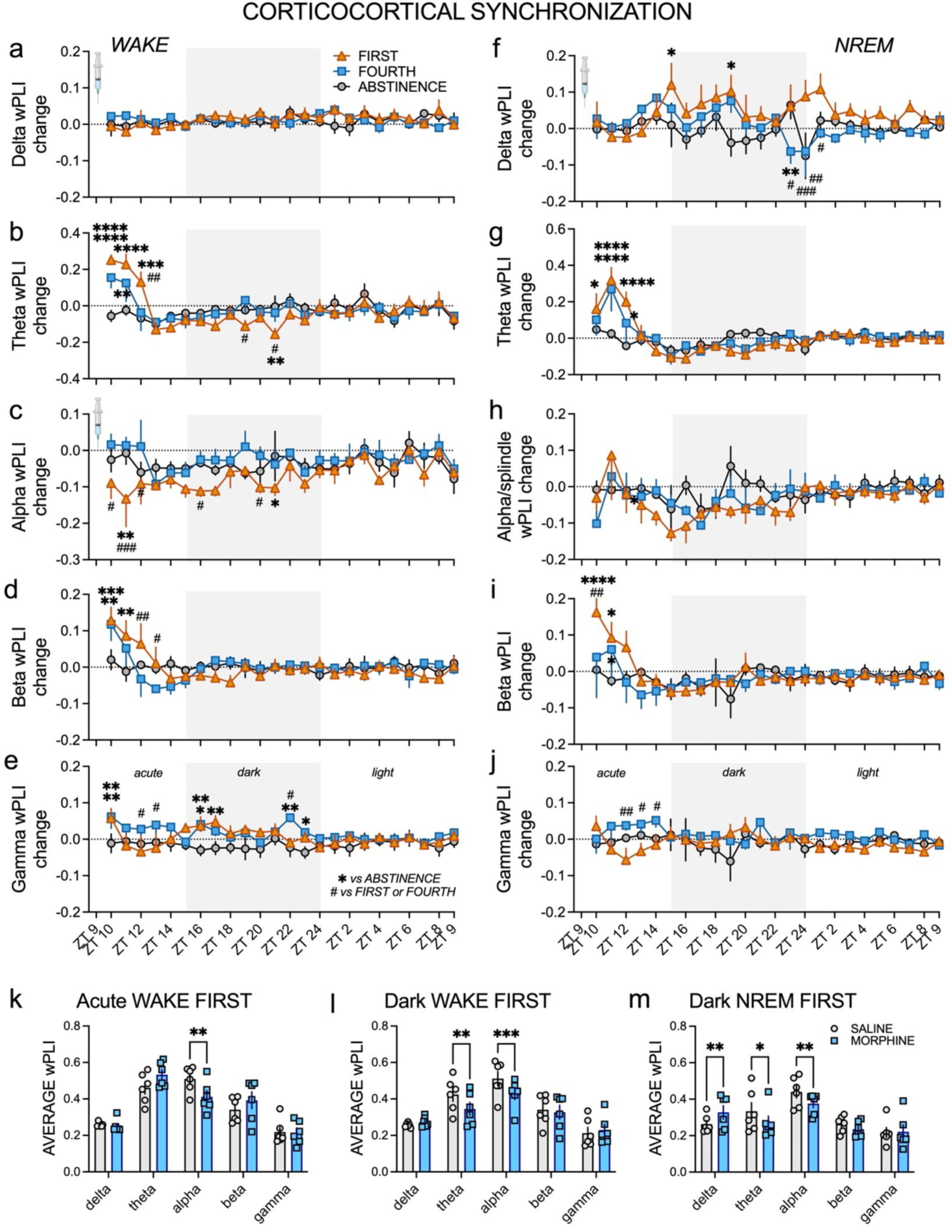
Corticocortical synchronization across WAKE and NREM following repeated morphine exposure and abstinence. Changes in cortico-cortical weighted phase-lag index (wPLI) are shown across recording time points for WAKE (a-e) and NREM (f-j) in the delta (a, f), theta (b, g), alpha (c, h), beta (d, i), and gamma (e, j) frequency bands. Data are presented for the first morphine exposure (orange triangles), fourth morphine injection (blue squares), and abstinence (gray circles). Values represent change in wPLI relative to the corresponding baseline/reference period; the horizontal dotted line at 0 indicates no change from baseline. Gray shading denotes the dark phase, with the subsequent unshaded region representing the light phase; the initial unshaded period corresponds to the acute post-treatment interval. Syringe symbols indicate treatment administration. Wake two-way RM ANOVA results: <u>Theta:</u> Time F_23,115_=5.86, p<0.0001, Condition F_2,10_=1.05, p=0.384, Interaction F_46,230_=2.82, p<0.0001; <u>Alpha:</u> Time F_23,115_=1.62, p=0.05, Condition F_2,10_=10.51, p=0.003, Interaction F_46,230_=0.98, p=0.515; <u>Beta</u>: Time F_23,115_=3.62, p<0.0001, Condition F_2,10_=0.006, p=0.993, Interaction F_46,230_=1.61, p=0.01; <u>Gamma</u>: Time F_23,115_=1.24, p=0.114, Condition F_2,10_=3.73, p=0.06, Interaction F_46,230_=1.26, p=0.140. NREM Mixed-effect model (REML) results for Condition: <u>Delta</u>: Time F_23,115_=1.08, p=0.378, Condition F_2,10_=3.65, p=0.064, Interaction F_46,211_=1.41, p=0.05; <u>Theta</u>: Time F_23,115_=7.05, p<0.0001, Condition F_2,10_=0.42, p=0.665, Interaction F_46,212_=1.94, p=0.0009; <u>Alpha:</u> Time F_23,115_=2.36, p=0.001, Condition F_2,10_=0.90, p=0.436, Interaction F_46,212_=0.88, p=0.694; <u>Beta</u>: Time F_23,115_=2.56, p=0.0005, Condition F_2,10_=0.27, p=0.769, Interaction F_46,212_=1.21, p=0.189; <u>Gamma</u>: Time F_23,115_=0.56, p=0.948, Condition F_2,10_=2.76, p=0.111, Interaction F_46,212_=1.26, p=0.143; Šídák’s post hoc was presented in the figures. (k–m) Average absolute cortico-cortical wPLI during selected intervals for saline (gray/white bars and circles) and morphine (blue bars and squares). (k) Average wPLI during the acute wake period following the <u>first exposure</u>; two-way RM ANOVA: Frequency F_4,20_=33.15, p<0.0001; Morphine F_1,5_ =0.05, p=0.835; Interaction F_4,20_=6.22, p=0.002, Šídák’s post hoc presented in the figure. (l) Average wPLI during dark phase wake episodes following the <u>first exposure</u>; two-way RM ANOVA: Frequency F_4,20_=16.29, p<0.0001; Morphine F_1,5_=6.99, p=0.046; Interaction F_4,20_=8.17, p=0.0004, Šídák’s post hoc presented in the figure. (m) Average wPLI during dark phase NREM following the <u>first exposure</u>: two-way RM ANOVA: Frequency F_4,20_=6.96, p=0.001; Morphine F_1,5_ =3.44, p=0.123; Interaction F_4,20_ =8.77, p=0.0003, Šídák’s post hoc presented in the figure. N=6 animals

In the averaged analyses, changes after the first injections were significant (Figure 6k–m), morphine reduced alpha synchronization during acute wakefulness (Figure 6k) and decreased theta and alpha synchronization during dark phase wakefulness (Figure 6l). During dark phase NREM sleep, morphine increased delta and reduced theta/alpha synchronization, with no differences in beta or gamma (Figure 6m). Alpha was therefore the most consistently affected band, differing during acute and dark phase wakefulness, as well as during dark phase NREM sleep.

### Thalamocortical synchronization

Thalamocortical synchronization also varied with frequency band, vigilance state, and exposure (Figure 7a-j; hourly analyses, wPLI differences from saline were shown and analyzed). The largest changes in thalamocortical synchronization occurred during the acute and dark phase periods. During wakefulness (Figure 7a-e), delta synchronization diverged early: it increased after the first injection and decreased after the fourth (Figure 7a). Theta synchronization rose transiently after the first injection, then decreased and remained reduced throughout the dark period; while after the fourth injection, a decrease appeared several hours before dark onset (Figure 7b). Alpha synchronization was predominantly reduced, without differing between conditions (Figure 7c), and beta decreased early after the fourth injection (Figure 7d). Gamma synchronization was reduced after both injections during the first half of the dark phase and elevated during abstinence (Figure 7e).

**Figure 7.**
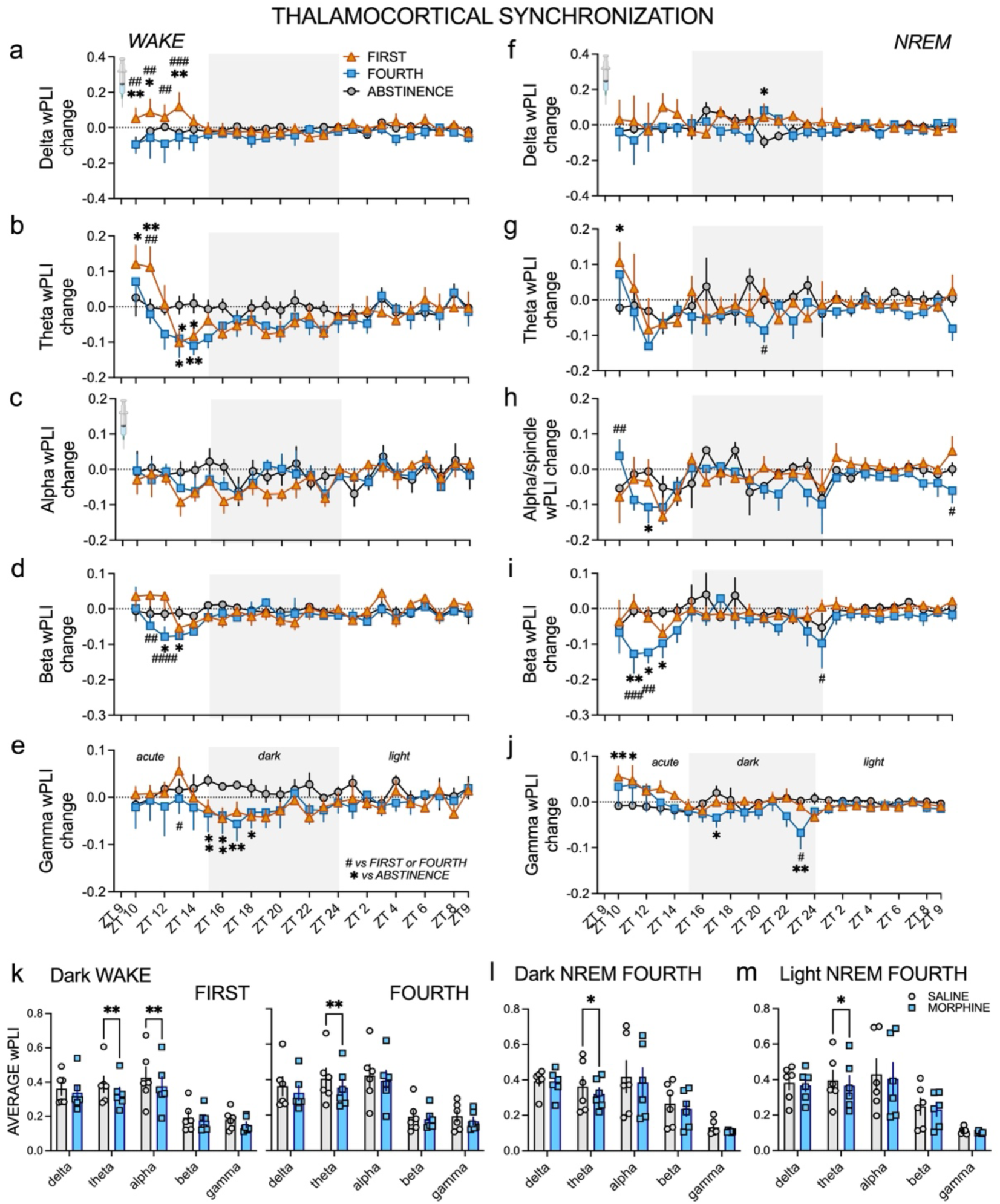
Thalamocortical synchronization across wakefulness and NREM sleep following repeated morphine exposure and abstinence. Changes in thalamocortical wPLI are shown across recording time points for wakefulness - wake (a-e) and NREM (f-j) in the delta (a, f), theta (b, g), alpha (c, h), beta (d, i), and gamma (e, j) frequency bands. Data are presented for the first morphine exposure (orange triangles), fourth morphine exposure (blue squares), and abstinence (gray circles). Values represent changes in wPLI relative to the corresponding baseline/reference period, with the horizontal dotted line at 0 indicating no change from baseline. Gray shading denotes the dark phase, with the subsequent unshaded region representing the light phase; the initial unshaded interval represents the acute post-treatment period. Syringe symbols indicate treatment administration. Wake two-way RM ANOVA results: <u>Delta:</u> Time F_23,115_=0.54, p=0.953, Condition F_2,10_=1.80, p=0.215, Interaction F_46,230_=1.21, p=0.178, <u>Theta:</u> Time F_23,115_=3.07, p<0.0001, Condition F_2,10_=1.17, p=0.351, Interaction F_46,230_=1.39, p=0.06; <u>Beta</u>: Time F_23,115_=1.52, p=0.08, Condition F_2,10_=1.44, p=0.282, Interaction F_46,230_=1.21, p=0.184; <u>Gamma</u>: Time F_23,115_=1.45, p=0.104, Condition F_2,10_=0.87, p=0.447, Interaction F_46,230_=1.08, p=0.353. NREM Mixed-effect model (REML) results for Condition: <u>Delta</u>: Time F_23,115_=0.22, p=0.999, Condition F_2,10_=0.87, p=0.449, Interaction F_46,212_=0.81, p=0.799; <u>Theta</u>: Time F_23,115_=1.09, p=0.366, Condition F_2,10_=1.84, p=0.209, Interaction F_46,212_=0.88, p=0.682; <u>Alpha:</u> Time F_23,115_=1.92, p=0.01, Condition F_2,10_=2.50, p=0.131, Interaction F_46,212_=1.27, p=0.135; <u>Beta</u>: Time F_23,115_=1.90, p=0.01, Condition F_2,10_=1.76, p=0.22, Interaction F_46,212_=1.31, p=0.101; <u>Gamma</u>: Time F_23,115_=2.37, p=0.001, Condition F_2,10_=0.72, p=0.508, Interaction F_46,212_=1.67, p=0.008; Šídák’s post hoc is presented in the figures. (k–m) Average absolute thalamocortical wPLI for saline (gray/white bars and circles) and morphine (blue bars and squares) during selected intervals. (k) Average wPLI during dark phase wake episodes following the first and fourth injections. During the <u>first exposure</u>, significant saline-morphine differences were observed in theta and alpha synchronization: two-way RM ANOVA: Frequency F_4,20_=9.30, p=0.0002; Morphine F_1,5_=6.41, p=0.05; Interaction F_4,20_=1.97, p=0.138, Šídák’s post hoc presented in the figure; during the <u>fourth injection</u>, a significant difference was observed in theta: two-way RM ANOVA: Frequency F_4,20_=7.81, p=0.0006; Morphine F_1,5_=2.22, p=0.196; Interaction F_4,20_ =6.22, p=0.002, Šídák’s post hoc presented in the figure. (l) Average wPLI during dark phase NREM following the <u>fourth exposure</u>, with a significant difference in theta synchronization: two-way RM ANOVA: Frequency F_4,20_=7.52, p=0.0007; Morphine F_1,5_=3.29, p=0.129; Interaction F_4,20_=0.698, p=0.602, Šídák’s post hoc presented in the figure. (m) Average wPLI during light phase NREM following the <u>fourth exposure</u>, with a significant difference in theta synchronization; two-way RM ANOVA: Frequency F_4,20_=7.85, p=0.0006; Morphine F_1,5_=4.13, p=0.098; Interaction F_4,20_=0.63, p=0.646, Šídák’s post hoc presented in the figure. N=6 animals

During NREM sleep (Figure 7f-j), delta synchronization fluctuated around baseline, with a transient dark phase reduction during abstinence (Figure 7f), and theta rose briefly after the first injection and decreased transiently during the dark phase after the fourth (Figure 7g). Alpha synchronization rose immediately after the fourth injection, then fell later in the acute response and at the end of the light phase (Figure 7h). Beta synchronization decreased markedly and acutely after the fourth injection, with a further decrease at the end of the dark phase (Figure 7i). Gamma synchronization rose modestly but persistently after the injections, then decreased later in the dark phase after the fourth (Figure 7j).

In the averaged analyses, morphine changed synchronization mainly after the repeated injections during the dark phase (Figure 7k-m). Morphine reduced theta synchronization during wakefulness after both exposures (Figure 7k left, right), and during NREM sleep after the fourth injection in both phases (Figure 7l,m). Alpha synchronization also decreased during dark phase wakefulness after the first injection (Figure 7k left). The remaining conditions were not significant. Alteration in theta frequency range was therefore the most persistent thalamocortical effect, present across exposures and in both states but confined to the dark period.

## DISCUSSION

To our knowledge, this is the first report of alterations in CMT oscillations and thalamocortical synchronization across repeated opioid exposure. Repeated morphine exposure reorganized sleep-wake regulation and oscillatory activity across cortical and thalamic networks in ways that depend on vigilance state, frequency band, recording site, time after administration, and exposure history. Two key findings are evident: the cortex and CMT often shift in opposite directions, indicating the thalamus is not merely a passive recipient of cortical sleep activity; additionally, the measures recover over different timescales during abstinence, making sleep quantity a poor indicator of sleep physiology. The wPLI further shows that morphine shifts network communication itself, not only the macroscopic composition of sleep.

### Sleep-wake architecture: morphine simultaneously increases locomotor activity and promotes wakefulness

The present findings align with an extensive literature on interactions between opioid signaling and sleep-wake regulation. Opioid receptors and endogenous opioid peptides are distributed throughout circuits governing arousal, sleep, respiration, and behavioral-state control^25^. Clinical reviews report that opioid exposure alters sleep-stage distribution and that chronic opioid therapy is associated with impaired sleep quality and excessive daytime sleepiness^4,26–28^. In healthy adults, single bedtime doses of sustained-release morphine or methadone significantly reduced SWS^26^. In our experiments, morphine markedly increased wakefulness, strongly suppressed NREM, and nearly abolished REM sleep (Figure 1). It also produced longer wake episodes, increased total wake duration, and substantial reductions in NREM sleep and, particularly, REM sleep (Figure 2). Morphine consistently increased the distance traveled and mean locomotor speed, while decreasing immobility. This effect did not diminish over repeated treatments (Figure 2), aligning with previously reported hyperlocomotion observed with moderate to high doses doses^29–31^. O’Brien et al. reported the same profile in mice and showed that opioid-induced wakefulness is electrophysiologically distinct from spontaneous wakefulness, describing the opioid condition as a dissociated state of consciousness^3^. The early wakefulness is therefore not an inability to initiate sleep, but an actively aroused yet electrophysiologically abnormal state. Repeated morphine injections did not produce an identical sleep pattern. After the fourth dose, acute wakefulness was promoted, and both NREM and REM sleep were suppressed. However, morphine-treated animals spent more time in NREM sleep than controls during the dark phase, followed by an increase in REM sleep (Figure 1). This can be explained by the accumulated sleep need that drives sleep recovery. This is in accordance with the two-process model^32^ and other reports showing that prior wake duration influences NREM SWA and sleep intensity^33–35^.

### Cortical and thalamic oscillatory activity

In the barrel cortex, morphine increased delta and low-gamma power, whereas alpha power decreased primarily after the first exposure (Table 1). Delta activity is conventionally associated with synchronized cortical activity during NREM sleep, yet McKelvey et al. showed that morphine generates slow EEG activity while rats remain behaviorally awake^28^. The increase observed here may therefore represent an abnormal cortical state in which slow-wave activity intrudes into pharmacologically maintained wakefulness. Cortical wake and NREM delta power peaked 4-5h after injection, rose more rapidly after the fourth than after the first injection, and remained elevated through both exposures and abstinence (Figure 3). This time course is consistent with a contribution of accumulated sleep pressure. Cortical theta power was transiently reduced and then moderately elevated in abstinence (Figure 3). Cortical alpha power decreased significantly only after the first exposure, indicating that the cortical alpha response adapts with repeated exposure. This differs from a study in rats, where cortical wake and NREM alpha power remained decreased during the initial days of exposure^28^. Cortical beta power showed a moderate, transient reduction during NREM sleep after the first injection, which recovered with repeated exposure and increased overall after repeated injections, but not during abstinence (Figure 3), again differing from the overall decrease reported in rats^28^. Low-gamma power increased immediately after morphine, more noticeably after the fourth injection, and acutely during NREM sleep, and remained moderately elevated across the dark and light phases and during abstinence (Figure 3). Previous findings indicate that gamma activity is typically linked to activated cortical states and that morphine affects gamma activity in other circuits^32^. So, rather than producing a uniform shift toward either sleep or arousal, morphine in our experiments generates a mixed electrophysiological state containing both slow-wave and higher-frequency activation, supporting opioid-induced dissociated state of consciousness^3^. Delta and gamma rise peaked at comparable latencies (4h and 3h, Figure 3), coinciding with the hyperlocomotion in the first 3-5h. It has been shown that locomotion itself increases gamma power in the nucleus accumbens as distance traveled increases^32^. The early spectral changes in cortical and central CMT observed here likely result from the combined effects of opioids directly impacting arousal circuits and the highly stimulated behavioral state caused by morphine. Also, it has been shown that morphine causes a distinctive cortical EEG pattern that includes morphine-induced spindles and high-voltage, low-frequency activity, with spectral features changing over the course of the drug effect^33^.

Direct involvement of the CMT is supported by the distribution of MOR within the thalamus. MOR is highly expressed in several intralaminar and midline thalamic nuclei, including the CMT, consistent with earlier autoradiographic evidence of μ-opioid binding throughout medial and midline thalamic regions^34,35^. Moreover, chronic morphine exposure produces adaptations in MOR signaling within medial thalamic neurons, including reduced opioid responsiveness in thalamic neurons and projection-dependent adaptations at terminals^36^. The acute and exposure-dependent changes in CMT oscillations observed here, hence, may arise, at least in part, from direct opioid modulation of CMT neuronal activity and from network-level effects mediated by other opioid-sensitive circuits.

The results from thalamic recordings reinforce both the distinction between the first and fourth exposures and the difference between cortex and CMT, with changes more pronounced during NREM sleep. During NREM sleep, thalamic delta power was initially reduced after the first injection, unchanged after the fourth injection, and slightly elevated during abstinence (Figure 4; Table 1). Across the dark and light phases, thalamic delta power rose substantially after both injections during wakefulness and NREM sleep (Figure 4). CMT theta and beta power were acutely suppressed after the first exposure and increased overall thereafter (Figure 4). Alpha power changed similarly to the cortex, whereas low-gamma power increased transiently during wake and NREM after the exposures, while in the cortex it remained elevated longer.

Our most critical finding is that delta power can be simultaneously elevated in the cortex and reduced in CMT, most evidently during the reduced NREM episodes that occur under morphine. The elevation of cortical NREM delta power was more pronounced after repeated exposure, whereas the reduction in CMT NREM delta power was less marked after the fourth injection (Figure 4). The CMT is well placed to generate such a dissociation because it participates in both wake promotion and NREM slow-wave organization. Tonic activation of CMT neurons induces NREM-to-wake transitions, whereas burst activation enhances cortical slow-wave synchronization^7^. Additionally, CMT activity contributes to sleep recovery after prolonged wakefulness^7,8^, and changes in CMT activity precede or accompany changes in neocortical activity during transitions into NREM sleep and anesthesia^6^. This implies that the same thalamic system can support both arousal and sleep-associated SWA, depending on its firing mode. These findings argue against treating the CMT as a passive recipient of cortical sleep activity.

### Relationship between vigilance-state occupancy and spectral power

Here we compared correlations using Fisher’s z transformation^37,38^. Our results show altered associations between state occupancy and spectral power, but do not establish causality. Morphine altered these associations differently in the cortex and CMT (Table 3). After the first exposure, cortical correlations with wake time changed significantly in the theta, alpha, and beta bands; while the correlations were positive under saline, they shifted toward zero or became negative under morphine. Thalamic wake correlations were unchanged after injection but increased in theta, alpha, and beta during abstinence, as did cortical wake-theta and wake-alpha correlations. During NREM sleep, cortical correlations with theta and alpha power increased and then decreased during abstinence, whereas thalamic correlations with delta and theta power increased and did not change during abstinence. Slope analysis showed a decreased cortical wake-theta slope and an increased cortical NREM-theta slope after injection (Table 2).

Two features stand out. First, coupling between NREM occupancy and CMT low-frequency power strengthened after the fourth exposure even though thalamic delta power was transiently reduced, indicating progressive engagement of the CMT-dependent mechanisms^7,39^. Second, during abstinence, cortical and CMT wake relationships became more positive across theta/alpha and beta bands, while cortical NREM relationships remained altered, strengthening negative relationships in theta/alpha bands.

### Corticocortical and thalamocortical synchronization

wPLI weights each phase-difference estimate by its magnitude and discounts zero-lag coupling, reducing sensitivity to volume conduction, shared sources, noise, and sample-size bias^18^. After the first exposure, corticocortical synchronization during wakefulness changed in a frequency- and time-dependent manner (Figure 6). Theta wPLI rose and then fell below baseline, with an overall dark phase decrease, and alpha wPLI decreased acutely, with mean acute wake alpha synchronization significantly lower under morphine. Beta synchronization shifted positively and was then suppressed, and low-gamma coupling rose transiently after injection and again at dark onset. During NREM sleep, dark phase delta synchronization increased while theta and alpha synchronization decreased as in wakefulness; a redistribution toward slower frequencies rather than a uniform change. The alpha result converges across all three levels of analysis: cortical alpha power decreased, wake-alpha relationships were disrupted, and corticocortical alpha synchronization decreased. Repeated administration delayed the acute rise in wake gamma wPLI and increased it later in the dark phase, reduced corticocortical NREM delta synchronization late in the dark phase, opposite to the first exposure, and transiently raised theta and beta wPLI (Figure 6e,f). During abstinence, synchronization returned largely to control values.

After the first exposure, thalamocortical delta wPLI increased during the early wake interval (Figure 7), while animals showed pronounced wakefulness and hyperlocomotion and thalamic delta power was reduced. Part of the sleep-associated thalamocortical network therefore engages while behavioral arousal is maintained, extending the demonstration of sleep-like slow cortical activity during opioid-induced wakefulness^28^. Theta synchronization changed most, rising early and then suppressed through the dark phase in both states. Wake alpha synchronization was disrupted acutely and remained reduced across the dark phase after the first exposure, whereas NREM alpha wPLI rose immediately after the fourth injection and then fell. Beta synchronization decreased transiently in both states after the fourth injection only. Low-gamma wake synchronization decreased at dark onset after both exposures, while NREM gamma synchronization remained elevated acutely and decreased by the end of the dark phase.

To our knowledge, wPLI has not previously been applied to opioid effects on sleep, although related work indicates that opioids reorganize oscillatory coordination. Morphine disrupts long-range gamma synchrony in hippocampal slices while leaving local oscillations intact^40^. It also narrows the spectral content of cortical activity in animal^33^ and human^41^ recordings, a "monorhythmization" toward a slower, more uniform state with anterior delta-theta predominance. While chronic exposure reduces dendritic branching and spine density in prefrontal and parietal cortex^42,43^, acute morphine perturbs brain-wide activity propagation until tolerance develop^44^, and fMRI during abstinence shows a durable signature of prior exposure on brain communication^45^. In our experiments spectral power and wPLI did not always change in parallel: increased cortical delta power did not invariably correspond to increased corticocortical delta synchronization, and increased gamma power coexisted with small or negative changes in gamma synchronization. Power quantifies magnitude at one site, wPLI phase consistency between sites, so morphine modifies amplitude and coordination independently.

#### Sleep pressure contributes to, but does not explain, the repeated-exposure phenotype

Whereas the first injection produced prominent changes in cortical alpha power and in the organization of wakefulness, increased cortical delta power emerged mainly after repeated exposure and persisted into abstinence. Repeated morphine-induced wakefulness offers a plausible mechanism for this transition. It is known that sleep pressure accumulates with time spent awake^46,47^. Additionally, experimental sleep deprivation in mice causes slow activity to emerge progressively during wakefulness^48^. NREM SWA increases during recovery sleep after deprivation^49^. Chronic or repeated sleep restriction produces persistent, regionally heterogeneous changes in cortical delta and faster-frequency activity^50^. In the closest anatomical parallel, a fast-delta component recorded in the barrel cortex and CMT, same recording sites as in our experiments, after prolonged wakefulness declined rapidly during recovery sleep and was specifically affected by manipulating CMT activity^39^.

Our findings support the existence of recurrent wakefulness, NREM rebound, increasingly prominent delta activity, and synchronizing coupling between NREM occupancy and thalamic low-frequency power^39,46–50^. Sleep pressure cannot, however, account for the entire phenotype since acute morphine itself generates slow waves during wakefulness^28^. Additionally, opioid-sensitive PVT neurons contribute directly to morphine-induced wakefulness^10^, and opioid withdrawal produces complex changes in NREM delta activity that need not resemble conventional recovery from sleep deprivation^51^. More importantly, the CMT showed reduced delta power immediately after the first exposure with no acute change after repeated injection. Increased delta activity should be interpreted as a homeostatic response to recurrent sleep disruption superimposed on direct opioid- and abstinence-dependent changes, rather than as evidence of increased sleep pressure.

### Recovery of sleep quantity during abstinence does not indicate complete physiological recovery

Cortical and thalamic spectral abnormalities and altered state-power relationships remained detectable during abstinence, whereas interregional phase synchronization returned to control values. Time spent awake or asleep can then be recovered before the electrophysiological organization of those states has normalized. This distinction is consistent with studies of spontaneous opioid withdrawal in which sleep architecture and EEG spectral characteristics follow different recovery trajectories. Persistent sleep and spectral abnormalities occur during spontaneous morphine withdrawal in mice, with withdrawal-associated NREM delta activity that does not increase uniformly^51^. Additionally, persistent sleep and quantitative EEG abnormalities occur across repeated spontaneous morphine withdrawal in rats^28^. The persistent spectral changes should also be considered in the context of altered nociceptive processing. Withdrawal from chronic morphine produces opioid withdrawal-induced hyperalgesia and mechanical hypersensitivity in mice^52–54^. Experimental nociceptive stimulation produces frequency-specific changes in cortical and thalamic electrophysiology, including enhanced low-frequency and gamma activity and suppression of alpha- and beta-range activity^55,56^, and the thalamus is a major node of ascending nociceptive processing^57^. Because persistent changes in delta, alpha, beta, and gamma activity coincided with mechanical hypersensitivity during abstinence, the abstinence EEG may reflect a composite post-opioid state in which sleep, arousal, and pain-related processes remain functionally coupled, rather than a residual signature of sleep deprivation alone. The contribution of the nociceptive state to these abnormalities remains to be tested experimentally.

### Methodological interpretation of wPLI and other limitations

Some methodological limitations may constrain interpretation. First, wPLI measures functional phase synchronization, not anatomical or directional connectivity^18^; an increase indicates a more consistent non-zero phase relationship but does not establish that the thalamus drives the cortex. Although CMT activity can precede cortical state transitions and influence cortical slow-wave organization^6,7,39^, more directed measures (Granger causality, phase-slope index) would be needed to test directionality. Second, translation to humans requires caution: rodents show polyphasic sleep and species-specific EEG characteristics, and human opioid exposure usually occurs alongside chronic pain, opioid use disorder, concomitant medications, and sleep-disordered breathing. Third, the acute post-injection interval and the subsequent phases are not equivalent: long-duration averages can obscure transient network responses, which is why we used time-resolved analysis. Fourth, the time-resolved spectral and wPLI analyses include only wakefulness and NREM sleep, because REM sleep was virtually absent in all morphine-treated animals during the acute intervals.

## CONCLUSION

Acute morphine administration produced hyperlocomotion and prolonged wakefulness that nonetheless exhibited sleep-associated electrophysiology: increased slow-wave activity, enhanced thalamocortical delta synchronization, while alpha power and coordination were disrupted. With repeated exposure, NREM and REM recovery became more prominent, CMT delta activity increased, and NREM occupancy coupled more strongly to low-frequency CMT activity. During abstinence, sleep architecture and interregional synchronization largely recovered while spectral and state-power abnormalities persisted alongside mechanical hypersensitivity. Because wPLI is comparatively insensitive to volume conduction, this redistribution is best interpreted as a genuine shift in network communication. It identifies the CMT-cortical network as a candidate substrate linking repeated opioid-induced wakefulness to altered NREM regulation, extending the role of the thalamus beyond PVT-mediated arousal mechanisms^10^. Morphine does not merely reduce sleep, it reorganizes it: cortex and CMT move in opposite directions, thalamocortical phase synchronization is redistributed, and the resulting spectral and state-power abnormalities outlast the recovery of sleep architecture itself.

## Funding

This study was funded in part by grants from the National Institutes of Health (GRANT# R35 GM141802 to S.M.T. and GRANT# K01 DA055258 to T.T.S)

## Acknowledgments

We thank the University of Colorado Anschutz Medical Campus Rodent In Vivo Neurophysiology Core for providing facilities to acquire video-EEG data. During the preparation of this work, the authors used ChatGPT-CU Anschutz (EDU) and Claude Science to assist with English-language polishing and sentence-structure refinement. After using these tools, the authors reviewed and edited the output for accuracy and clarity, and they take full responsibility for the overall integrity of the final publication.

## Conflict of interest

The authors received no compensation and have no conflicting financial interests related to the work described in this manuscript.

## Notes

### Competing Interest Statement

The authors have declared no competing interest.

