## Supplemental for "Repeated morphine reorganizes sleep-wake states, cortical and central medial thalamic oscillations, and network synchronization in mice"

Table S1. Direction of morphine-induced change in spectral power during the dark phase (10h). Each cell gives the first injection/fourth injection/abstinence, in that order. ↑ statistically significant increase, ↓ statistically significant decrease, — no significant change reported.

| | Cortex Wake | Cortex NREM | Cortex REM | CMT Wake | CMT NREM | CMT REM |
|---|---|---|---|---|---|---|
| **Delta** | ↑↑↑ | ↑↑↑ | ↑—— | ↑↑— | ↑↑— | |
| **Theta** | ↑↑↑ | ↑↑↑ | ↑—— | ↑↑— | ↑↑— | |
| **Alpha** | —↑— | ↑↑ — | | —↑— | ↑↑— | —↑↓ |
| **Beta** | ↑↑— | ↑↑ — | | —↑— | ↑↑— | |
| **Gamma** | —↑↑ | ↑—— | ↑—— | | | |

Table S2. Direction of morphine-induced change in spectral power during the light phase (9h). Each cell gives the first injection/fourth injection/abstinence, in that order. ↑ statistically significant increase, ↓ statistically significant decrease, — no significant change reported.

| | Cortex Wake | Cortex NREM | Cortex REM | CMT Wake | CMT NREM | CMT REM |
|---|---|---|---|---|---|---|
| **Delta** | ↑↑↑ | ↑↑↑ | —↑— | ↑↑— | ↑↑— | ↑↑— |
| **Theta** | ↑↑— | ↑↑— | | ↑↑— | ↑↑— | ↑—— |
| **Alpha** | —↑— | ↑↑— | | —↑— | ↑↑— | ↑—— |
| **Beta** | —↑— | ↑↑— | —↑— | —↑— | ↑—— | |
| **Gamma** | —↑↑ | —↑↑ | —↑— | | ↑—— | |

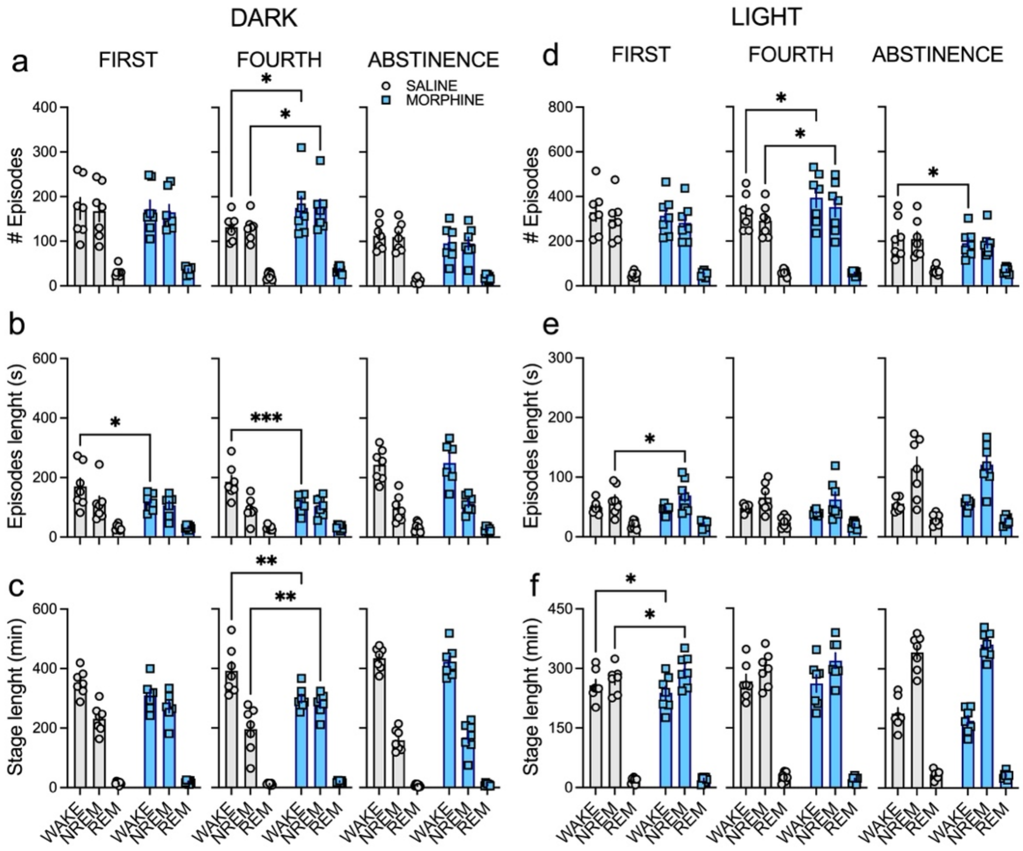


**Figure S1. Morphine-induced alterations in sleep-wake architecture during repeated exposure and abstinence during the dark and light phases.** (a, b) Number of vigilance-state episodes recorded during the first injection, fourth injection, and abstinence period during dark (a) and light (b) phase. Saline-treated animals are shown as open circles/gray bars, whereas morphine-treated animals are shown as blue squares/bars. Morphine significantly altered episode frequency, with more pronounced effects during repeated exposure during the light phase. (c, d) Mean duration of vigilance-state episodes during dark (c) and light (d) phase across recording sessions. Morphine modified episode length in state- and time-dependent manners, particularly during initial and repeated exposures. (e, f) Total time spent in wakefulness (wake), non-rapid eye movement sleep (NREM), and rapid eye movement sleep (REM) during dark (e) and light (f) phase after the first/fourth injections and abstinence. Morphine shifted the distribution of vigilance states by increasing wakefulness and reducing NREM sleep during treatment, with partial normalization during abstinence. Bars represent mean ± SEM with individual data points overlaid. Statistical significance is denoted by asterisks (*p<0.05, **p<0.01, ***p<0.001).

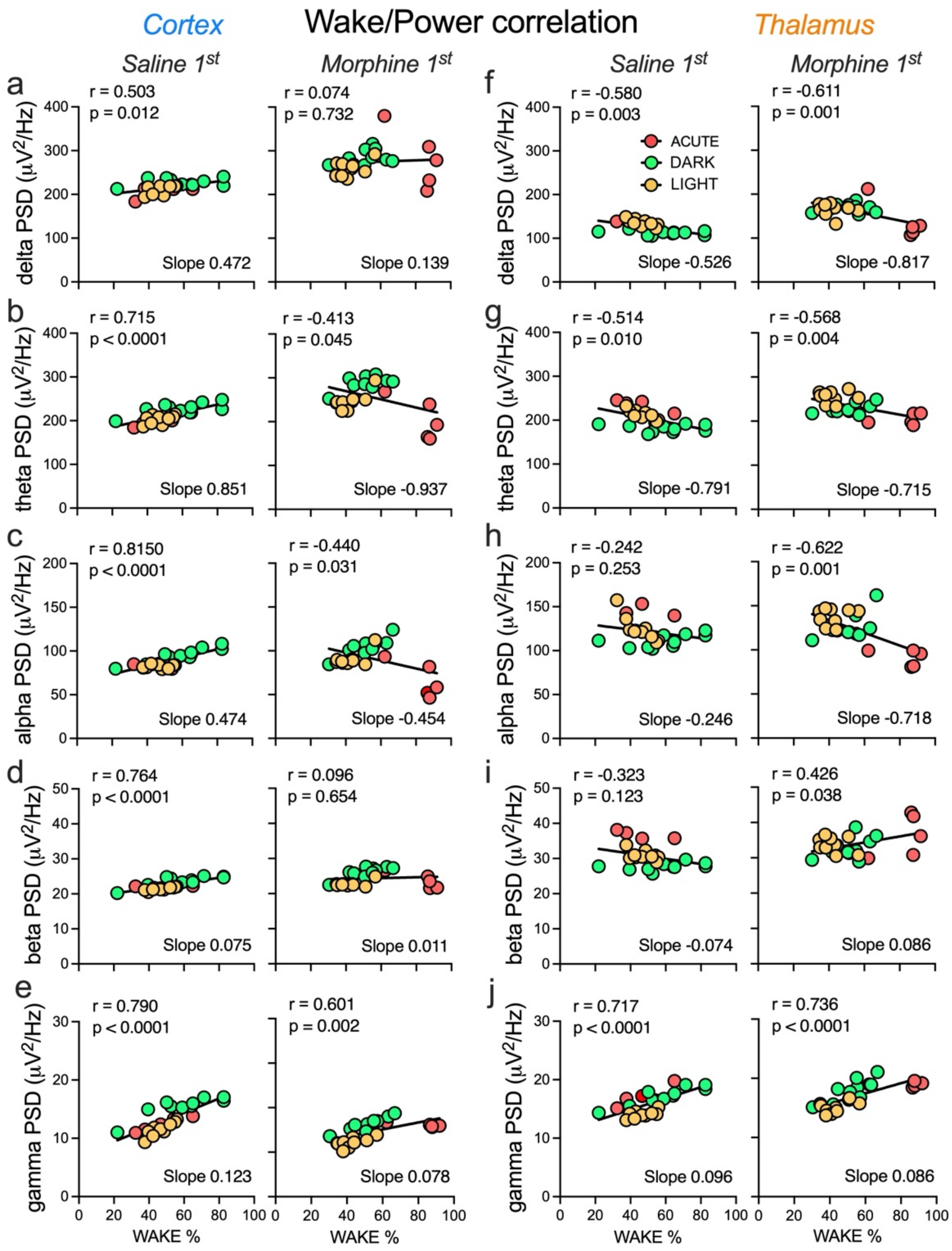


**Figure S2. Correlation between wakefulness and oscillatory power in cortical and thalamic recordings following FIRST saline and morphine administration.** Scatterplots illustrate the relationship between the percentage of time spent awake (wake %) and spectral power density (PSD; μV²/Hz) within individual frequency bands. Data are shown separately for cortical (a-e) and thalamic (f-j) recordings following the first saline administration and the first morphine administration. Points represent individual recordings obtained during the acute (red), dark phase (green), and light phase (yellow) conditions. Black lines indicate linear regression fits. Pearson correlation coefficients (r), significance values (p), and regression slopes are shown in each panel. Cortex: (a) Delta-band power versus wakefulness, (b) Theta-band power versus wakefulness, (c)

Alpha-band power versus wakefulness, (d) Beta-band power versus wakefulness, (e) Gamma-band power versus wakefulness. Thalamus: (f) Delta-band power versus wakefulness, (g) Theta-band power versus wakefulness, (h) Alpha-band power versus wakefulness, (i) Beta-band power versus wakefulness, (j) Gamma-band power versus wakefulness.

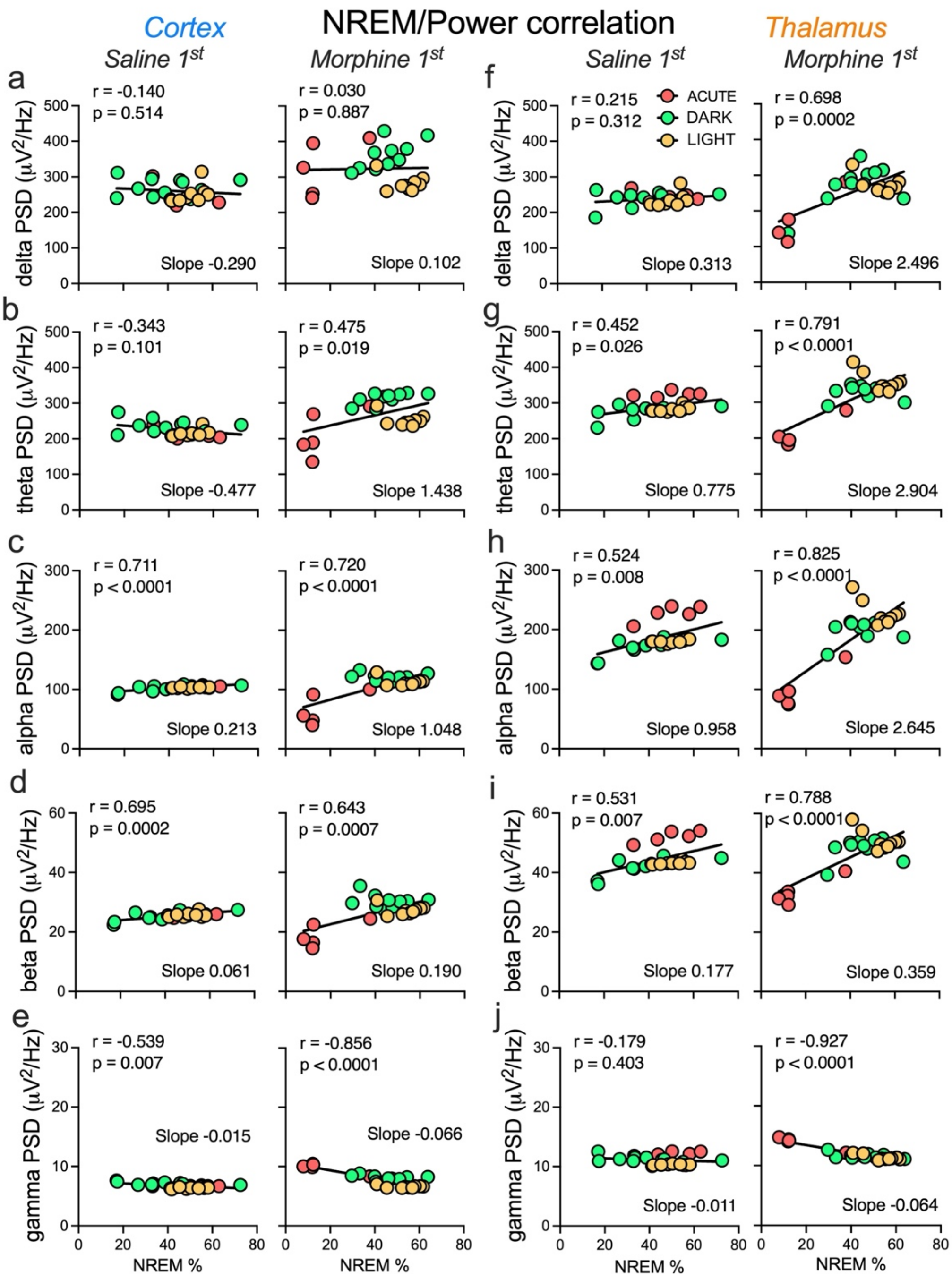


**Figure S3. Correlation between NREM and oscillatory power in cortical and thalamic recordings following FIRST saline and morphine administration.** Scatterplots illustrate the relationship between the percentage of time spent in NREM sleep (NREM %) and spectral power density (PSD; μV²/Hz) within individual frequency bands. Data are shown separately for cortical (a-e) and thalamic (f-j) recordings following the first saline administration and the first morphine administration. Points represent individual recordings obtained during the acute (red), dark phase (green), and light phase (yellow) conditions. Black lines indicate linear regression fits. Pearson correlation coefficients (r), significance values (p), and regression slopes are shown in each panel. Cortex: (a) Delta-band power versus wakefulness, (b) Theta-band power versus wakefulness, (c) Alpha-band power versus wakefulness, (d) Beta-band power versus wakefulness, (e) Gamma-band power versus wakefulness. Thalamus: (f) Delta-band power versus wakefulness, (g) Theta-band power versus wakefulness, (h) Alpha-band power versus wakefulness, (i) Beta-band power versus wakefulness, (j) Gamma-band power versus wakefulness.

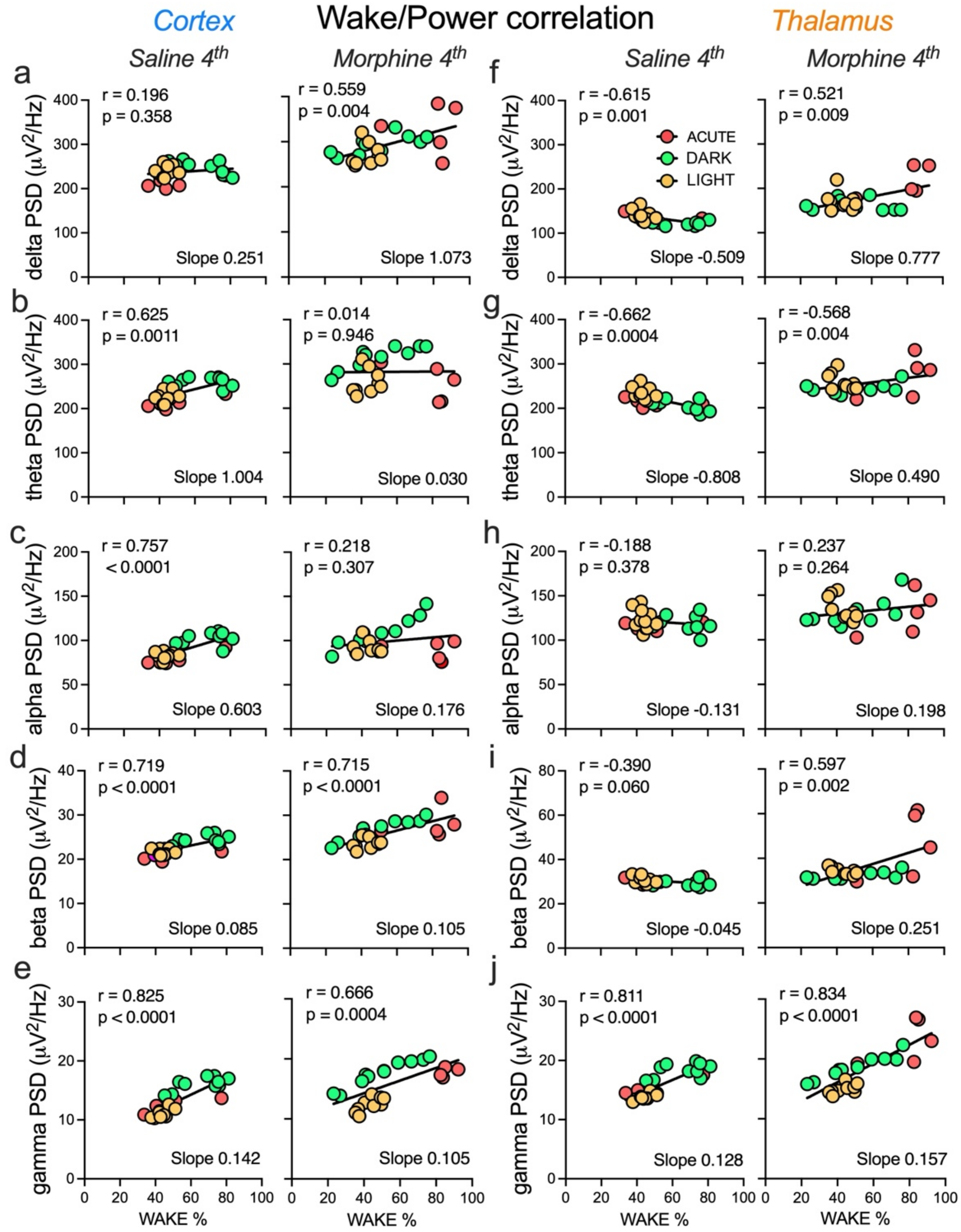


**Figure S4. Correlation between wakefulness and oscillatory power in cortical and thalamic recordings following FOURTH saline and morphine administration.** Scatterplots illustrate the relationship between the percentage of time spent awake (wake %) and spectral power density (PSD; μV²/Hz) within individual frequency bands. Data are shown separately for cortical (a-e) and thalamic (f-j) recordings following the fourth saline administration and the fourth morphine administration. Points represent individual recordings obtained during the acute (red), dark phase (green), and light phase (yellow) conditions. Black lines indicate linear regression fits. Pearson correlation coefficients (r), significance values (p), and regression slopes are shown in each panel. Cortex: (a) Delta-band power versus wakefulness, (b) Theta-band power versus wakefulness, (c) Alpha-band power versus wakefulness, (d) Beta-band power versus wakefulness, (e) Gamma-band power versus wakefulness. Thalamus: (f) Delta-band power versus wakefulness, (g) Theta-band power versus wakefulness, (h) Alpha-band power versus wakefulness, (i) Beta-band power versus wakefulness, (j) Gamma-band power versus wakefulness.

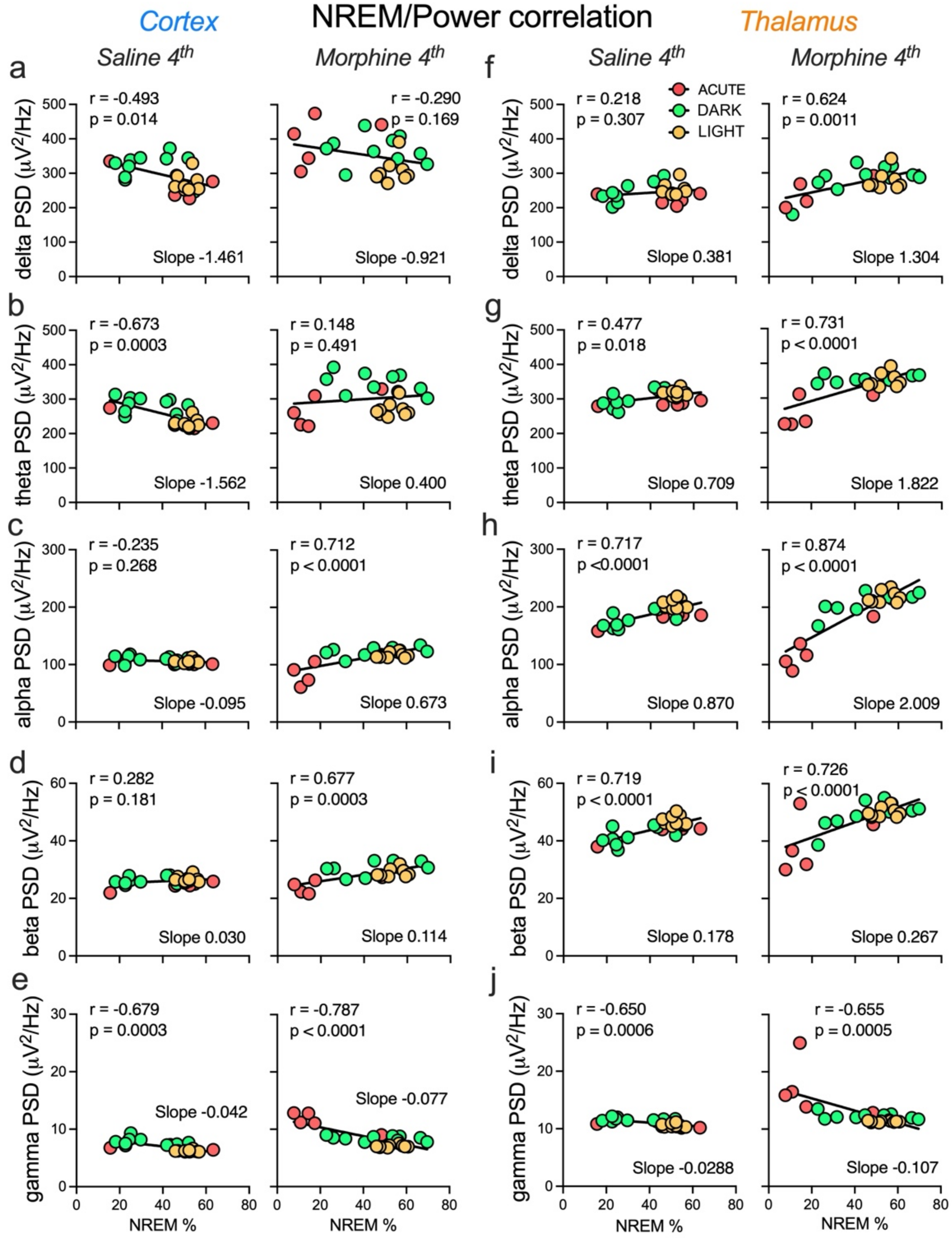


**Figure S5. Correlation between NREM and oscillatory power in cortical and thalamic recordings following FOURTH saline and morphine administration.** Scatterplots illustrate the relationship between the percentage of time spent in NREM sleep (NREM %) and spectral power density (PSD; μV²/Hz) within individual frequency bands. Data are shown separately for cortical (a-e) and thalamic (f-j) recordings following the fourth saline administration and the fourth morphine administration. Points represent individual recordings obtained during the acute (red), dark phase (green), and light phase (yellow) conditions. Black lines indicate linear regression fits. Pearson correlation coefficients (r), significance values (p), and regression slopes are shown in each panel. Cortex: (a) Delta-band power versus wakefulness, (b) Theta-band power versus wakefulness, (c) Alpha-band power versus wakefulness, (d) Beta-band power versus wakefulness, (e) Gamma-band power versus wakefulness. Thalamus: (f) Delta-band power versus wakefulness, (g) Theta-band power versus wakefulness, (h) Alpha-band power versus wakefulness, (i) Beta-band power versus wakefulness, (j) Gamma-band power versus wakefulness.

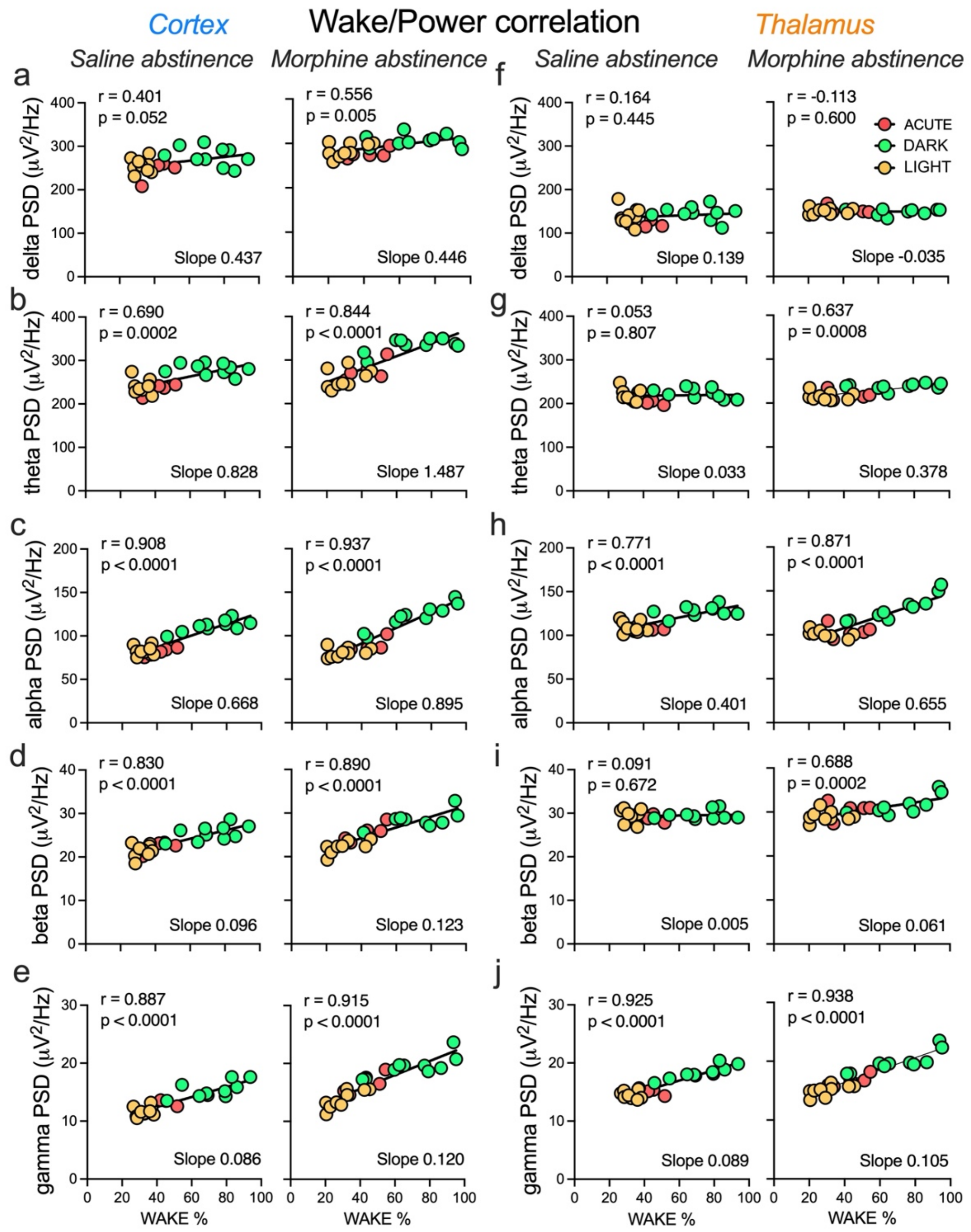


**Figure S6. Correlation between wakefulness and oscillatory power in cortical and thalamic recordings following ABSTINENCE.** Scatterplots illustrate the relationship between the percentage of time spent awake (wake %) and spectral power density (PSD; μV²/Hz) within individual frequency bands. Data are shown separately for cortical (a-e) and thalamic (f-j) recordings following the saline and morphine abstinence. Points represent individual recordings obtained during the acute (red), dark phase (green), and light phase (yellow) conditions. Black lines indicate linear regression fits. Pearson correlation coefficients (r), significance values (p), and regression slopes are shown in each panel. Cortex: (a) Delta-band power versus wakefulness, (b) Theta-band power versus wakefulness, (c) Alpha-band power versus wakefulness, (d) Beta-band power versus wakefulness, (e) Gamma-band power versus wakefulness. Thalamus: (f) Delta-band power versus wakefulness, (g) Theta-band power versus wakefulness, (h) Alpha-band power versus wakefulness, (i) Beta-band power versus wakefulness, (j) Gamma-band power versus wakefulness.

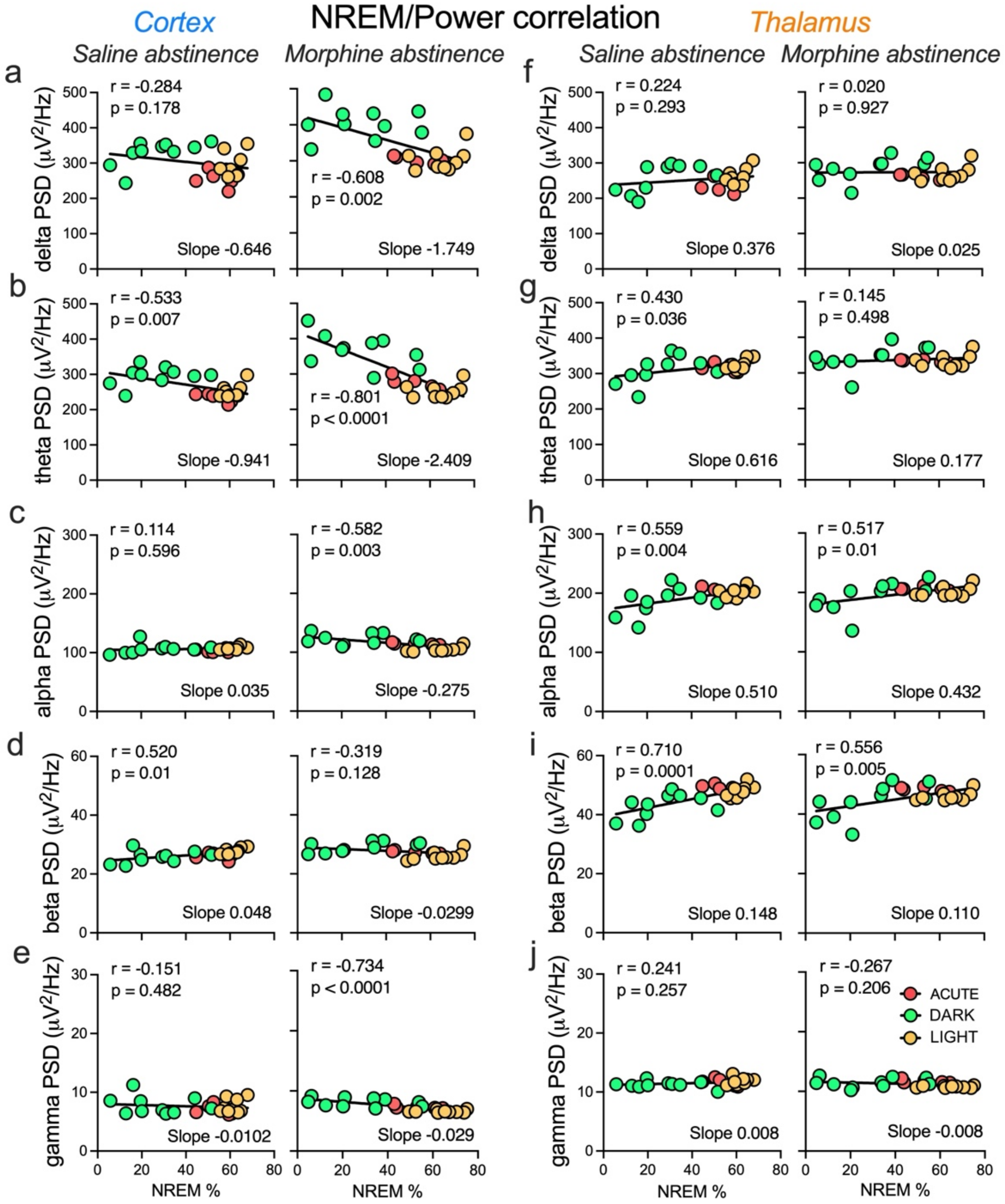


**Figure S7. Correlation between NREM and oscillatory power in cortical and thalamic recordings following ABSTINENCE.** Scatterplots illustrate the relationship between the percentage of time spent in NREM sleep (NREM %) and spectral power density (PSD; μV²/Hz) within individual frequency bands. Data are shown separately for cortical (a-e) and thalamic (f-j) recordings following saline and morphine abstinence. Points represent individual recordings obtained during the acute (red), dark phase (green), and light phase (yellow) conditions. Black lines indicate linear regression fits. Pearson correlation coefficients (r), significance values (p), and regression slopes are shown in each panel. Cortex: (a) Delta-band power versus wakefulness, (b) Theta-band power versus wakefulness, (c) Alpha-band power versus wakefulness, (d) Beta-band power versus wakefulness, (e) Gamma-band power versus wakefulness. Thalamus: (f) Delta-band power versus wakefulness, (g) Theta-band power versus wakefulness, (h) Alpha-band power versus wakefulness, (i) Beta-band power versus wakefulness, (j) Gamma-band power versus wakefulness.
